# Mechanisms for dysregulation of excitatory-inhibitory balance underlying allodynia in dorsal horn neural subcircuits

**DOI:** 10.1101/2024.06.10.598179

**Authors:** Alexander G. Ginsberg, Scott F. Lempka, Bo Duan, Victoria Booth, Jennifer Crodelle

## Abstract

Chronic pain is a wide-spread condition that is debilitating and expensive to manage, costing the United States alone around $600 billion in 2010. In a common type of chronic pain called allodynia, non-painful stimuli produce painful responses with highly variable presentations across individuals. While the specific mechanisms remain unclear, allodynia is hypothesized to be caused by the dysregulation of excitatory-inhibitory (E-I) balance in pain-processing neural circuitry in the dorsal horn of the spinal cord. In this work, we analyze biophysically-motivated subcircuit structures that represent common motifs in neural circuits in layers I-II of the dorsal horn. These circuits are hypothesized to be part of the neural pathways that mediate two different types of allodynia: static and dynamic. We use neural firing rate models to describe the activity of populations of excitatory and inhibitory interneurons within each subcircuit. By accounting for experimentally-observed responses under healthy conditions, we specify model parameters defining populations of subcircuits that yield typical behavior under normal conditions. Then, we implement a sensitivity analysis approach to identify the mechanisms most likely to cause allodynia-producing dysregulation of the subcircuit’s E-I signaling. We find that disruption of E-I balance generally occurs either due to downregulation of inhibitory signaling so that excitatory neurons are “released” from inhibitory control, or due to upregulation of excitatory neuron responses so that excitatory neurons “escape” their inhibitory control. Which of these mechanisms is most likely to occur, the subcircuit components involved in the mechanism, and the proportion of subcircuits exhibiting the mechanism can vary depending on the subcircuit structure. These results suggest specific hypotheses about diverse mechanisms that may be most likely responsible for allodynia, thus offering predictions for the high interindividual variability observed in allodynia and identifying targets for further experimental studies on the underlying mechanisms of this chronic pain condition.

**Author summary:** While chronic pain affects roughly 20% of the US adult population [1], symptoms and presentations of the condition are highly variable across individuals and its causes remain largely unknown. A prevailing hypothesis for the cause of a type of chronic pain called allodynia is that the balance between excitatory and inhibitory signaling pathways between neuron populations in the spinal cord dorsal horn may be disrupted. To help better understand neural mechanisms underlying allodynia, we analyze biologically-motivated mathematical models of subcircuits of neuron populations that are part of the pain processing signaling pathway in the dorsal horn of the spinal cord. We use a novel sensitivity analysis approach to identify mechanisms of subcircuit dysregulation that may contribute to two different types of allodynia. The model results identify specific subcircuit components that are most likely to contribute to each type of allodynia. These mechanisms suggest targets for further experimental study, as well as for pharmacological intervention for better pain treatments.

## 1 Introduction

Understanding the neural circuitry in the spinal cord that processes pain signals is vital for understanding the mechanisms responsible for chronic pain [2, 3], a wide-spread condition affecting ∼20% of adults in the US [1] and costing the US alone around $600 billion in 2010 [4]. Indeed, the spinal cord is responsible for the initial processing of both tactile and pain-inducing stimuli at the periphery, and for relaying them to the brain [5–7]. In particular, tactile and pain-inducing signals travel from the periphery to the spinal cord along different classes of afferent nerve fibers: *Aβ*-fibers which respond to innocuous stimuli such as gentle pressure or the brush of clothing on skin and *C*-fibers, and to some extent *Aδ* fibers, which respond to heat, noxious chemicals, or intense mechanical stimuli [5–7]. Signals associated with painful stimuli, upon arriving at the spinal cord, are filtered by intermediate neural circuitry in the superficial laminae (primarily laminae I and II) of the spinal cord’s dorsal horn [8]. From there, the intensity of the painful signal is relayed to the brain through the firing activity of excitatory projection neurons in lamina I [8].

It is widely accepted that processing of afferent signals in the healthy pain-processing circuit of the dorsal horn relies on a balance between excitation and inhibition [2, 8, 9]. Melzack and Wall originally proposed this hypothesis in 1965 as the conceptual “gate control” model [10], and aspects of the theory remain influential and relevant [8, 9, 11]. Namely, the idea that inhibitory neurons “gate” the activity of excitatory neurons which relay pain signals towards the brain [2] remains a key hypothesis. In particular, while both excitatory and inhibitory neural populations receive input from *Aβ* fibers, inhibitory neurons suppress the firing of the excitatory neurons, thus blocking the response of projection neurons to pain-inducing signals from *C*-fibers.

Pathological changes to dorsal horn neural circuitry is frequently proposed as the culprit behind chronic pain [12, 13], including allodynia, a type of chronic pain in which individuals feel pain in response to normally innocuous stimuli [14]. There are several types of allodynia classified according to the type of innocuous stimuli that is painful [14]. In this work, we focus on two well-studied types of allodynia: static and dynamic [15, 16]. In static allodynia, individuals feel pain in response to gentle pressure that normally would not be painful [16], while in dynamic allodynia individuals feel pain in response to brushing-type stimuli that normally would also not be painful [16].

Recent experimental results indicate that different types of excitatory and inhibitory dorsal horn neurons mediate static and dynamic allodynia, as discussed in Section 1.1. Specifically, rodent experiments have shown that either static or dynamic allodynia can be induced by activating or inactivating (e.g. ablating) specific populations of excitatory or inhibitory interneurons in the dorsal horn [16–19], creating a disruption of excitatory-inhibitory (E-I) balance. In clinical conditions, however, allodynia likely occurs through more subtle circuit disruptions. Since E-I balance in a circuit can be achieved by diverse contributions of different excitatory and inhibitory neural populations, its disruption leading to allodynia can potentially occur through multiple pathways.

In this work, we use biophysically-based mathematical modeling to identify likely mechanisms by which dorsal horn neural subcircuits may be dysregulated to produce allodynia. While both A*β* fiber and C fiber activity play a role in gate control [20] and thus in allodynia, we mainly focus on the role of A*β* fiber activity in allodynia. In particular, we construct models of neural subcircuits implicated in mediating static and dynamic allodynia whose parameters are constrained to reproduce experimentally-observed behaviors. These constraints result in distributions of parameter sets representing populations of subcircuits that achieve E-I balance in different ways. We then identify the most sensitive mechanisms that disrupt E-I balance to result in allodynia in the subcircuit population. We find that the particular means of disruption varies across the subcircuit population and that the most sensitive mechanisms depend on subcircuit structure, thus predicting diverse, multiple mechanisms that may be most likely responsible for allodynia.

### 1.1 Proposed dorsal horn subcircuits mediating allodynia

Here, we describe subcircuit motifs that reflect recent experimental evidence for the structure of dorsal horn layer I-II networks mediating static allodynia (as evoked by a von Frey device) and dynamic allodynia (Fig 1A). Experimental studies in rodents have shown that static allodynia is reliant on activity in three putatively different types of excitatory interneurons: somatostatin-positive (SOM+) [16], calretinin-positive (CR+) [19], and protein kinase C *γ*-positive (PKC*γ*+) [17] cells. Additionally, inactivation of either dynorphin-positive (DYN+) [16] or parvalbumin-positive (PV+) [17] inhibitory interneurons is sufficient to produce static allodynia, suggesting that these inhibitory cells usually gate activation of SOM+, CR+ and PKC*γ*+ excitatory cells. Experiments have also identified direct synaptic connections from PV+ to PKC*γ*+ neurons [17].

**Fig 1.**
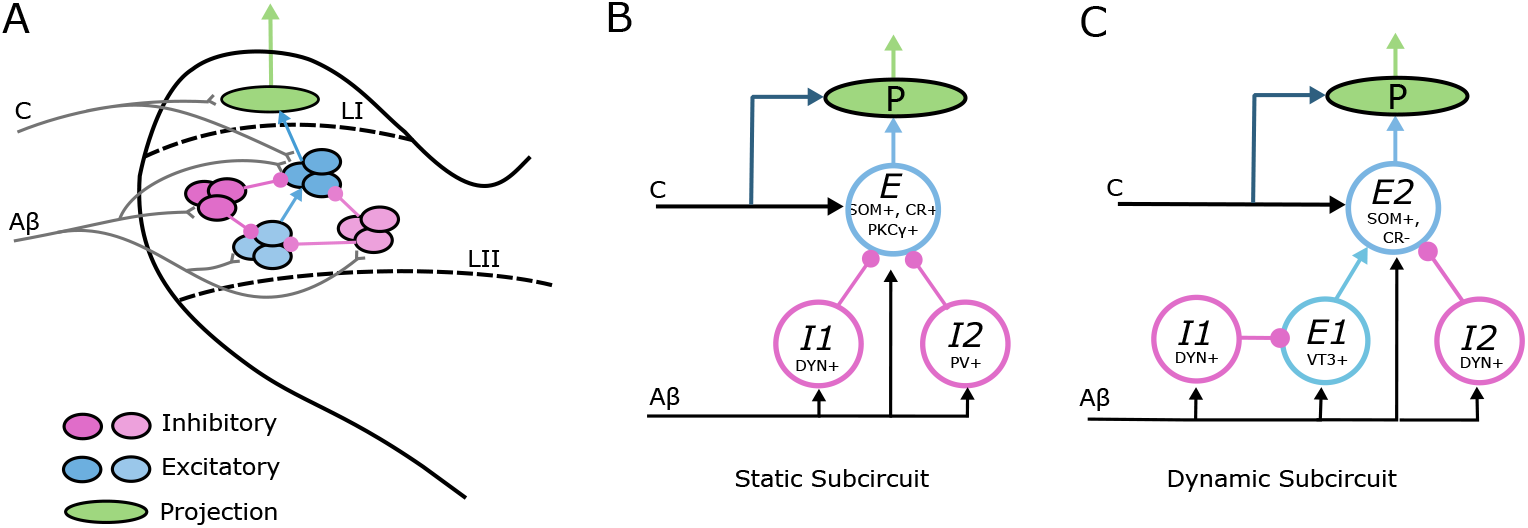
Proposed subcircuits in layer I-II of the dorsal horn mediating static and dynamic allodynia. **(A)** Schematic of neural circuitry in layer I-II of the dorsal horn consisting of populations of inhibitory (magenta shades) and excitatory (blue shades) interneurons that filter A*β* and C fiber inputs to projection neurons (green) that transmit signals to the brain. **(B)** A schematic of the proposed subcircuit mediating static allodynia. *I*1 and *I*2 represent the populations of dynorphin-positive (DYN+) and parvalbumin-positive (PV+) inhibitory interneurons, respectively, and *E* represents a collective population of somatostatin-positive (SOM+), calretinin-positive (CR+) and protein kinase C *γ*-positive (PKC*γ*+) excitatory interneurons. **(C)** A schematic of the proposed subcircuit mediating dynamic allodynia. *I*1 and *I*2 represent populations of DYN+ inhibitory neurons, and *E*1 and *E*2 represent the populations of VGLUT3-positive (VT3) and somatostatin-positive/calretinin-negative (SOM+/CR-) excitatory interneurons. In all panels, *Aβ* and *C* represent inputs relayed from the periphery along *Aβ* and *C* fibers, respectively.

Based on these experimental results, we consider a simplified subcircuit motif representing part of the pathway mediating static allodynia that consists of three neural populations (Fig 1B). One population, *E*, represents the collective activity of layer I-II SOM+, CR+ and PKC*γ*+ excitatory interneurons. The *E* population is inhibited by two distinct inhibitory interneuron populations, *I*1 representing DYN+ cells and *I*2 representing PV+ cells. All three populations receive input from *Aβ* fibers, (as suggested in [16, 19, 21]), while the *E* population is presumed to be additionally targeted by *C* fiber input (see e.g. [16, 19]).

Rodent experimental studies probing the mechanisms for dynamic allodynia have identified that it relies on the activity of SOM+ excitatory interneurons that do not express the calbindin 2/calretinin gene (CR-). Specifically, ablating SOM+ neurons abolishes or greatly reduces dynamic allodynia in mice [18]. On the other hand, ablating neurons expressing the calbindin 2/calretinin gene (CR+) had no effect on dynamic allodynia [18]. Thus, the SOM+ neurons that are necessary for dynamic allodynia must also be negative for calretinin (SOM+/CR-). Excitatory interneurons expressing vesicular glutamate transporter 3 (VT3+) also contribute to dynamic allodynia since their ablation or silencing eliminates or attenuates dynamic allodynia induced by nerve injury or ablation in mice [18]. Moreover, it has been proposed that the VT3+ neurons synapse onto the excitatory SOM+/CR-neurons [18]. Further experimental results suggest that activity of the VT3+ excitatory interneurons are gated by DYN+ inhibitory neurons since VT3+ neurons fire action potentials in response to *Aβ* input when inhibitory signaling is blocked [18]. Ablation of DYN+ cells is sufficient to induce dynamic allodynia [16] suggesting that DYN+ inhibitory cells may provide local inhibitory control of these two excitatory interneuron populations involved in dynamic allodynia.

To account for these experimental results, we consider a simplified subcircuit representing part of the pathway mediating dynamic allodynia that consists of four neural populations (Fig 1C). An excitatory population *E2* representing the collective activity of SOM+/CR-excitatory neurons is excited by a population *E1* representing VT3+ neurons. *E1* and *E2* receive inhibitory input from populations *I1* and *I2*, respectively, representing DYN+ cells. All populations receive *Aβ* input, (as suggested in [16, 18]), and *E2* is presumed to be additionally targeted by *C* fiber input (see [18]).

The *E* population in the static allodynia subcircuit and the *E2* population in the dynamic allodynia subcircuit are assumed to provide direct synaptic input to layer I projection cells that transmit painful signals to the brain [22]. Hence, painful stimuli generating activity on C fibers would activate these excitatory cells and the downstream projection neurons. Under normal healthy conditions, responses of these excitatory populations to non-painful stimuli signaled by *Aβ* input is assumed to be gated by the inhibitory populations. Allodynia occurs when the E-I balance is disrupted such that non-painful *Aβ* input causes firing in these excitatory populations, leading to activation of the downstream projection neurons. While allodynia can also be induced by nociceptor-mediated sensitization of projection neurons [20], here we focus on allodynia caused by disruption of inhibitory gating of A*β* input.

### 1.2 Overview

The goal of this study is to identify and characterize “most likely” potential mechanisms that disrupt the processing of A*β* signals and E-I balance in these subcircuits leading to allodynia. We model the activity and interactions of the neural populations in these subcircuits using a population firing rate model formalism (see e.g. [23–25]) constrained by experimental data. We begin by identifying the full space of model parameters that generate responses to A*β* signals that is correlated to healthy conditions in each subcircuit, i.e. low firing of the *E* or *E2* populations. Due to the relative mathematical simplicity of our firing rate model formalism, we are able to analytically define an “allowable parameter space” (APS) as a solution to a system of inequalities on model parameters. The APS for each subcircuit indicates the considerable range of parameter combinations that can reproduce healthy responses and demonstrates the myriad ways that E-I balance might be obtained. The population of subcircuits corresponding to the points in the APS represents the high variability in subcircuit structure that may occur physiologically.

Next, for each subcircuit, we define the analytic condition on model parameters that generates a response to A*β* input correlated with allodynia, i.e. high firing of the *E* or *E2* populations. Solving this condition defines a hypersurface in the model parameter space, which we call the “allodynia surface”, that separates parameter sets for which the subcircuit generates healthy, allodynia-free responses for all typical *Aβ* inputs, vs responses where allodynia may be present, (for at least some typical level of *Aβ* input). For model parameter sets (points) in the APS, we can then identify the minimal change in parameters that leads to allodynia by computing the shortest vector in parameter space from the point in the APS to the allodynia surface. The direction of these shortest vectors, corresponding to changes in specific model parameters, indicates the least variation of the subcircuit structure that results in the allodynia response. Since larger variations in parameter values can also generate an allodynia response, these shortest vectors represent the “most likely” potential mechanisms for allodynia to occur.

We then analyze the shortest vectors from the APS to the allodynia surface to identify distinct variations of parameters that dysregulate E-I balance leading to allodynia. Specifically, we find that the APS points form clusters based on parameters requiring the relatively smallest changes to reach the allodynia surface. These different clusters can be interpreted as representing different underlying mechanisms for generating allodynia.

We find that these proposed allodynia mechanisms generally involve the dysregulation of E-I balance occurring due to the release of excitatory cells from inhibitory gating or to the excitatory cells escaping the inhibitory gating. Significantly, the specific subcircuit components that are associated with the “release” or “escape” mechanisms differ in the subcircuits due to their differing network structures. As such, our results identify the diverse ways by which excitatory and inhibitory components within the subcircuits combine and interact to maintain E-I balance, and characterize the sensitivity of these subcircuits to disruptions that can lead to pathological responses.

The paper is organized as follows: in Section 2.1, in order to illustrate our analysis method, we first analyze a simple subcircuit representing the canonical gate control model. In Sections 2.2 and 2.3, we then apply the analysis to the proposed subcircuits mediating static and dynamic allodynia, respectively. In Section 3, we summarize the results and put them into a greater physiological context. Details of the models and analysis methods are contained in Section 4.

## 2 Results

For our models of layer I-II dorsal horn neuronal subcircuits, we use a firing-rate model formalism [23–25] that describes the average membrane voltage *V*_*x*_ (in mV) and average firing rate *f*_*x*_ (*x* = *E* or *I* or *x* = *Ej, Ij* for *j* = 1, 2; in Hz) of the populations of excitatory and inhibitory interneurons (see Section 4). In this formalism, average voltages are governed by equations of the form:

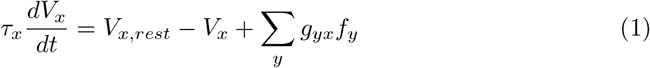

where *V*_*x,rest*_ is the average resting voltage (in mV) and *τ*_*x*_ is the time constant (in s) for the average voltage response of neurons in population *x*. Average population firing rates are computed by 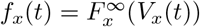 where the steady state firing rate activation function 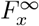 has a sigmoidal shape and its parameters are fit to experimental measurements of frequency-voltage relationships in dorsal horn neurons [26]. Input to the subcircuits from A*β* fibers is modeled by a compound Poisson process with piecewise-constant rate, (see Section 4.3 for details). To fit subcircuit responses to normal healthy conditions and to identify mechanisms for allodynia, we vary the synaptic coupling strengths between populations and from the A*β* fibers, *g*_*yx*_ (*y* = *E, I, Ej, Ij* or A*β*; in V-s). In particular, the APS for each subcircuit is a subset of the space of these parameters and the allodynia surface is a hypersurface defined within this space.

### 2.1 Analysis of a simple “gate control” subcircuit

We first illustrate our analysis methodology by applying it to a simple subcircuit representing the canonical “gate control” model (Fig 2A). This simple subcircuit consists of an inhibitory interneuron population (*I*) that inhibits an excitatory interneuron population (*E*). Both populations receive input from *Aβ* fibers. We assume that the *E* population directly synapses onto projection neurons, thus any sustained activity of the *E* population is a proxy for a painful response. Under normal healthy conditions, *Aβ* activity should not cause firing of the *E* population, i.e. a painful response should not occur. This is achieved through appropriate balancing of the inhibitory input from the *I* population and the excitatory A*β* input onto the *E* population.

**Fig 2.**
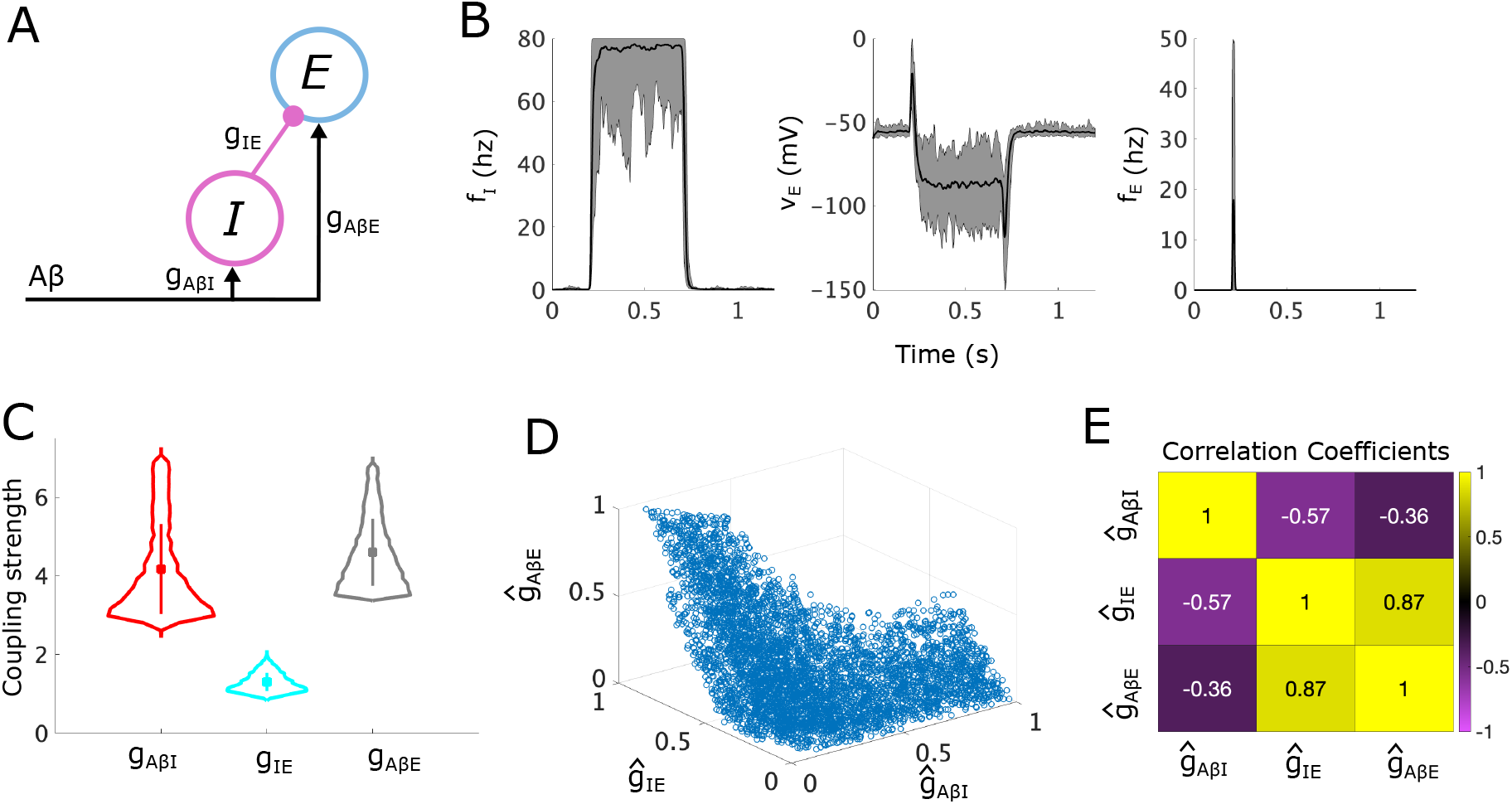
Allowable parameter space (APS) for the simple “gate control” subcircuit. **(A)** A schematic of the simple “gate control” subcircuit, where *E* and *I* represent populations of excitatory and inhibitory interneurons, respectively. **(B)** Simulation results showing the mean (black lines) and range (shaded gray areas) for the firing-rate responses of the *I* population (left panel), and the average voltage (middle panel) and firing rate (right panel) of the *E* population, simulated for 20 sampled points in the APS each with a different random *Aβ* stimulus in range [10, 20] Hz (during *t* ∈ [0.2, 0.7] s). **(C)** Violin plots showing the distribution of each coupling strength parameter with mean (square marker) and range of values that lie within one standard deviation (vertical bar) indicated. **(D)** A scatter plot of 5000 uniformly sampled APS points in the normalized (*ĝ*_*AβI*_, *ĝ*_*IE*_, *ĝ*_*AβE*_) space. **(E)** Normalized Pearson correlation coefficients between coupling strength parameters in the normalized APS sample.

For our analysis of this simple subcircuit, we first derive the conditions on the coupling strength parameters in the subcircuit, namely (*g*_*AβI*_, *g*_*IE*_, *g*_*AβE*_), that generate normal healthy responses to A*β* inputs and define the APS. We then mathematically express the allodynia surface *S* that separates (*g*_*AβI*_, *g*_*IE*_, *g*_*AβE*_) space into regions where the subcircuit displays healthy vs allodynia responses. Conveniently, we can visualize the APS and the allodynia surface *S* in the 3D parameter space, making this simple subcircuit especially helpful for illustrating our analysis method. Computing the shortest vectors in (*g*_*AβI*_, *g*_*IE*_, *g*_*AβE*_) space from the parameter points in the APS to the allodynia surface *S* identifies the minimal parameter changes that can induce an allodynia response. By clustering the parameter points in the APS by their shortest path vectors, we characterize the different ways that the coupling strengths between subcircuit components maintain and disrupt E-I balance in the subcircuit, thus suggesting underlying mechanisms responsible for allodynia.

#### The APS for the simple subcircuit

To constrain the subcircuit coupling strength parameters, we impose conditions on the steady-state voltages of the *E* and *I* populations in response to constant A*β* input to account for experimental observations. To start, we require that steady-state voltages of the *E* and *I* populations, 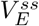 and 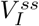, respectively, remain within biologically reasonable bounds, *V*_*min*_ and *V*_*max*_, in response to a sustained, typical non-painful input on the *Aβ* fibers. Throughout this study, we assume that non-painful input corresponds to *Aβ* fiber firing rate *f*_*Aβ*_ ∈ [10, 20] Hz as observed in slowly adapting mechanoreceptors in response to ramp-and-hold mechanical stimulation [27]. To incorporate the phenomenon of A*β* input gating responses to C fiber input, also known as pain inhibition, we require that the net signaling to the *E* population in response to typical *Aβ* activity is inhibitory, causing the voltage of the *E* population to decrease. To enforce this gating response under normal healthy conditions, we require that the *I* population fires for all typical, non-painful *f*_*Aβ*_ values, but the *E* population does not, thus representing a non-painful response. We also incorporate conditions accounting for experimental results showing that ablation of DYN+ inhibitory interneurons causes allodynia [16] by requiring that the *E* population fires in response to typical *Aβ* input in the absence of *I* population activity.

Each of these conditions leads to an inequality that steady-state voltages 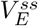 and 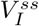 must satisfy (Table 1). These inequalities are derived by requiring the steady state solution of the average population voltage equation (Eq. 1) in response to a constant *f*_*Aβ*_ input remains below the maximum average voltage *V*_*x,max*_, above the minimum average voltage *V*_*x,min*_, above the voltage threshold for firing *V*_*x,thr*_, or below the resting voltage *V*_*x,rest*_ for the pain inhibition condition (*x* = *E, I*; see Section 4.2 for values of these bounds).

**Table 1.** Conditions and resulting inequalities used to constrain the simple subcircuit and define its allowable parameter space (APS). Condition type and Condition (first 2 columns) describe the rationale behind each condition. The resulting inequality on population steady-state voltages 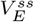 and 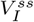 are given in the 3rd column. These inequalities must hold for typical non-painful *f*_*Aβ*_ values, which we take to be *f*_*Aβ*_ ∈ [10, 20] Hz. *V*_*x,max*_ and *V*_*x,min*_ are the maximum and minimum limits on average voltage, respectively, *V*_*x,rest*_ is the resting membrane voltage and *V*_*x,thr*_ is the voltage threshold for firing (*x* = *E, I*).

| Condition Type | Condition | Steady state voltage inequality |
| --- | --- | --- |
| Control conditions | $V_I$ upper bound | $V_{I,max} \geq V_I^{ss} = g_{A\beta I} f_{A\beta} + V_{I,rest}$ |
| | $I$ fires | $V_{I,thr} \leq V_I^{ss} = g_{A\beta I} f_{A\beta} + V_{I,rest}$ |
| | Pain inhibition | $V_{E,rest} \geq V_E^{ss} = g_{A\beta E} f_{A\beta} - g_{IE} f_I + V_{E,rest}$ |
| | $V_E$ lower bound | $V_{E,min} \leq V_E^{ss} = g_{A\beta E} f_{A\beta} - g_{IE} f_I + V_{E,rest}$ |
| $I$ ablation conditions | $V_E$ upper bound | $V_{E,max} \geq V_E^{ss} = g_{A\beta E} f_{A\beta} + V_{E,rest}$ |
| | $E$ fires | $V_{E,thr} \leq V_E^{ss} = g_{A\beta E} f_{A\beta} + V_{E,rest}$ |

These conditions can be re-written as a system of 6 inequalities, some nonlinear, on the coupling strengths (*g*_*AβI*_, *g*_*IE*_ and *g*_*AβE*_). The set of all coupling strength 3-tuples, (*g*_*AβI*_, *g*_*IE*_, *g*_*AβE*_), that satisfy these inequalities constitutes the APS for this subcircuit. In particular, subcircuit instantiations in the APS do not produce allodynia for typical non-painful *f*_*Aβ*_ signals (i.e. when *f*_*Aβ*_ ∈ [10, 20] Hz). Because these inequalities must be satisfied for a range of input *f*_*Aβ*_ signals, solving them explicitly requires finding the solution to a system of nonlinear optimization problems. As described in Section 4.4.1, this system for the simple subcircuit can be explicitly solved using Lambert functions. For the static and dynamic allodynia subcircuits, however, we do not find an explicit solution, although Lambert functions are used to greatly simplify the system of inequalities (Sections 4.4.2 and 4.4.3). Instead of explicitly solving this system of optimization problems, we compute a uniformly-distributed sample of the APS. Because the APS may be non-convex (see Fig 2D), we must develop a sampling algorithm (Section 4.6) to generate the uniform sample shown in Fig 2C - Fig 2D.

To illustrate that the points within the APS correspond to subcircuit instantiations which yield the desired behaviors, we simulate instances of the simple subcircuit model for 20 randomly sampled APS points in response to a noisy *Aβ* signal with mean firing rate in [10, 20] Hz. The subcircuit responses show that the firing rate of the *I* population rises to at least 80% of its maximum of 80 Hz (Fig 2B, left panel) and the inhibition from the *I* population causes voltages of the *E* population to drop (Fig 2B, middle panel), as we require for pain inhibition. Thus, as expected, *E* population firing rates remain low except for a brief increase at the onset of the A*β* stimulus (Fig 2B, right panel). On the other hand, if we simulate ablation of the *I* population (by setting *g*_*IE*_ to zero), the voltage of the *E* population sufficiently increases so that the *E* population fires in response to non-painful *Aβ* stimuli, thus simulating allodynia, (see S1 Fig in the supplementary material).

From the distributions of the coupling strength values (*g*_*AβI*_, *g*_*IE*_ and *g*_*AβE*_) across the APS (Fig 2C), we see that *g*_*AβI*_ has the largest range of allowable values (from about 2.6 - 7.1 mV*/*Hz), with *g*_*AβE*_ having the second largest range (about 3.5 - 6.9 mV*/*Hz), and *g*_*IE*_ having the smallest range (about 0.9 - 2.1 mV*/*Hz). Because *g*_*AβI*_ can thus be changed the most without exiting the APS, whereas *g*_*IE*_ can be changed the least, they are in some sense the least and most sensitive coupling strengths, respectively, maintaining E-I balance in this subcircuit.

To begin to understand mechanisms that can induce allodynia in terms of relative changes in *g*_*yx*_ (*y, x* = *E, I, Aβ*), we consider the normalized coupling strength as a proportion of its observed range over the APS, *ĝ*_*yx*_, and continue our analysis using the normalized APS in (*ĝ*_*AβI*_, *ĝ*_*IE*_, *ĝ*_*AβE*_) space. By normalizing parameters in this way, not only do we control for the effects of different ranges for each parameter, but we can better compare the effects of changing different parameters. In Fig 2D, we show the full set of normalized sampled APS points.

Considering the relationship among the coupling strengths in the APS can help identify mechanisms that contribute to the maintenance and eventual disruption of E-I balance in the subcircuit. The correlations between normalized coupling strengths (see Fig 2E) indicate that *ĝ*_*IE*_ and *ĝ*_*AβE*_ values are strongly positively correlated. (Note that correlations between normalized coupling strengths imply correlations between unnormalized coupling strengths, although the magnitude of the correlations may differ for the normalized vs unnormalized coupling strengths). This highlights that under normal conditions the direct excitatory and inhibitory inputs to the *E* population are balanced to maintain little-to-no firing in the *E* population. On the other hand, *ĝ*_*IE*_ and *ĝ*_*AβI*_ are negatively correlated indicating that inhibitory signaling to the *E* population (influenced by *ĝ*_*IE*_) may be conserved for varying responses of the *I* population to A*β* input (influenced by *ĝ*_*AβI*_). The slight negative correlation between *ĝ*_*AβE*_ and *ĝ*_*AβI*_ also suggests a balance in the subcircuit of excitatory and inhibitory responses to A*β* signaling.

#### Mechanisms for generating allodynia in the simple subcircuit

To determine the vulnerabilities of the simple subcircuit to allodynia, we find which changes in the coupling strengths most easily result in *E* population firing in response to non-painful *f*_*Aβ*_ input (in the range [10, 20] Hz). In our model formalism, allodynia is presumed to occur when the steady-state average voltage of the *E* population, 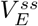, exceeds the firing threshold for some typical non-painful, sustained *f*_*Aβ*_:

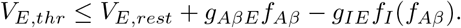

We use this inequality to define the allodynia surface *S* in (*g*_*AβI*_, *g*_*IE*_, *g*_*AβE*_) space, above which the corresponding subcircuit instantiation produces allodynia for at least one value of *f*_*Aβ*_ ∈ [10, 20] Hz. We use “above” in the sense that the *g*_*AβE*_ component of a point in parameter space defines a height for that point. This surface, *S*, is then the following set of points:

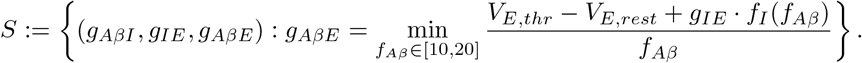

We compute *S* by solving this minimization problem in the unnormalized parameter space (see Section 4.7 for details). Plotting *S* in the normalized (*ĝ*_*AβI*_, *ĝ*_*IE*_, *ĝ*_*AβE*_) space shows that it always lies above the APS (Fig 3A).

**Fig 3.**
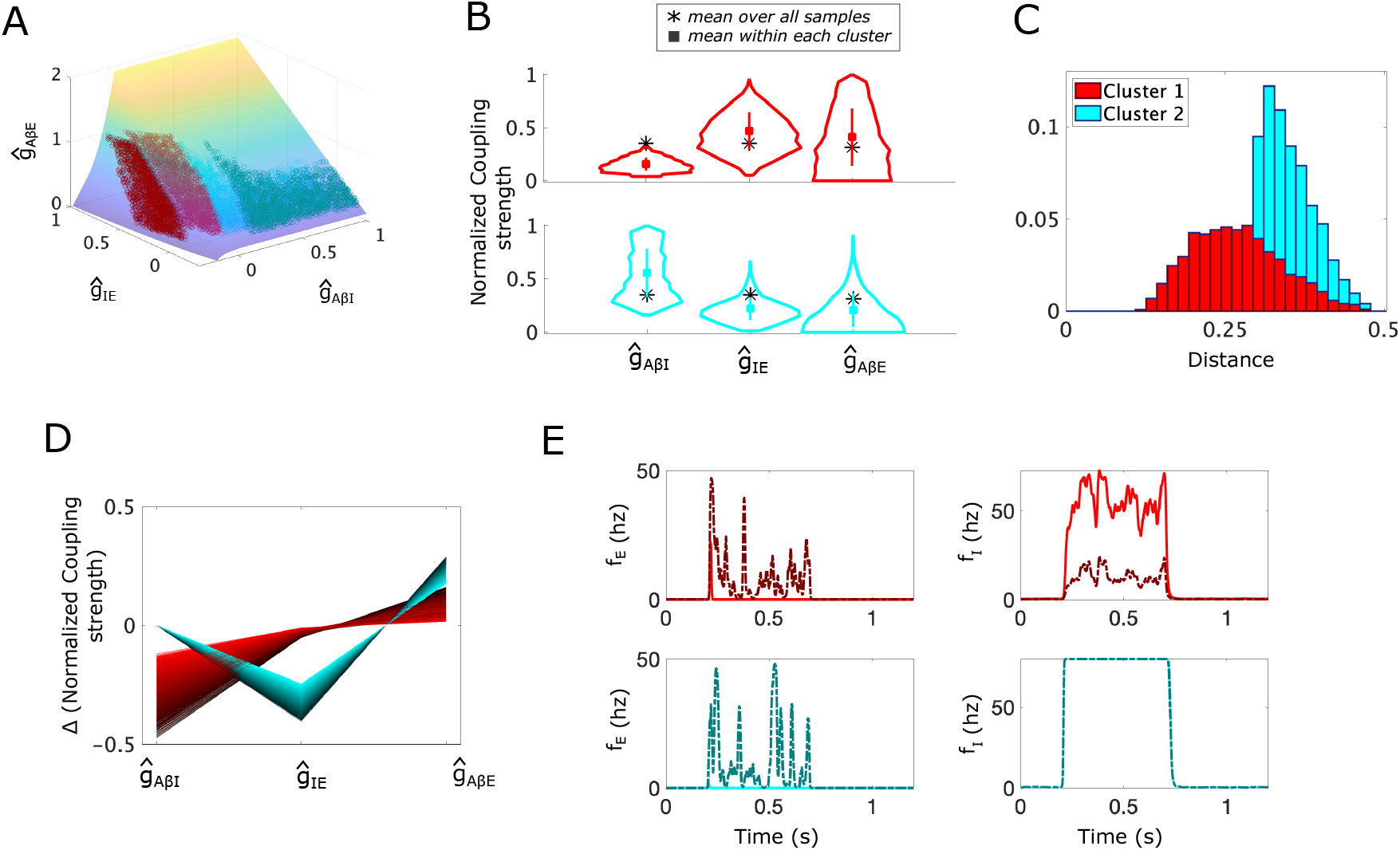
Allodynia mechanisms represented as shortest paths from the APS to the allodynia surface in the simple “gate control” subcircuit. **(A)** A scatter plot of the sampled APS points (5000 points) in the normalized (*ĝ*_*AβI*_, *ĝ*_*IE*_, *ĝ*_*AβE*_) space with the allodynia surface *S* overlaid. Based on the direction of their shortest paths, APS points separate into 2 clusters (Cluster 1 (red) and Cluster 2 (blue)). The nearest point on the allodynia surface from each sampled APS point is also shown (darker red and blue points on *S*). **(B)** Violin plots of the coupling strength distributions for the APS points in each cluster. Black ∗ shows the mean APS values, colored square shows the mean values in each cluster. **(C)** The probability distribution of the shortest distances to the allodynia surface (overall profile). The shading represents the contributions from each cluster to the overall profile. For instance, a bar that is 70% light blue indicates that cluster 2 constitutes 70% of subcircuit instantiations with the corresponding distance to the allodynia surface. **(D)** Parallel plot representation of the components of the shortest path vectors from APS points to their corresponding nearest points on *S*, colored according to cluster membership. **(E)** Firing rate responses *f*_*E*_ (left panels) and *f*_*I*_ (right panels) to a noisy *Aβ* input signal with amplitude *f*_*Aβ*_ ∈ [10, 20] Hz occurring during *t* ∈ [0.2, 0.7]s for each cluster, (top row: Cluster 1, bottom row: Cluster 2). Solid lines correspond to the subcircuit with coupling strength values set to the mean values for the cluster and dash-dotted lines correspond to the simple subcircuit instantiation with coupling strengths set to the corresponding closest point on the allodynia surface. *Aβ* input mean frequencies are chosen as the smallest value that induces allodynia for each cluster.

For each subcircuit instantiation associated with a point in the APS, we identify its vulnerability to allodynia by computing the shortest path from that point to the allodynia surface *S* using a customized global optimization scheme (see Section 4.8). Because the shortest path indicates how to reach the allodynia surface by altering coupling strengths as little as possible, it represents the direction in parameter space in which the subcircuit instantiation is most vulnerable to allodynia. In this way, the components of the shortest path vector suggest which subcircuit components will need to change, along with the corresponding magnitudes of their relative change, in order to disrupt *E* -*I* balance and induce allodynia.

To identify differences in vulnerabilities to allodynia (measured as changes in the parameters to reach the allodynia surface along the shortest path), we assign each point in the APS to a cluster based on the components of its shortest-path vector. The clustering algorithm, using density-based scanning [28], identifies two clusters in the APS for the simple subcircuit (Fig 3A). For points in Cluster 1 (red), the shortest direction to the allodynia surface primarily involves decreasing *ĝ*_*AβI*_, while for points in Cluster 2 (cyan), the shortest path involves decreasing *ĝ*_*IE*_ and increasing *ĝ*_*AβE*_.

Breaking down the points in each cluster, we find that points in Cluster 1 have larger values of *ĝ*_*IE*_ and *ĝ*_*AβE*_ and smaller values of *ĝ*_*AβI*_ relative to the mean of each coupling strength in the whole APS (Fig 3B). We also see that points in Cluster 1 are generally closer to the allodynia surface than Cluster 2 (Fig 3C), indicating that points in Cluster 1 are more sensitive to dysregulation than points in Cluster 2. In terms of the mechanism for allodynia, the shortest paths from Cluster 1 points to *S* consist of a decrease in *ĝ*_*AβI*_ and an increase in *ĝ*_*AβE*_ (Fig 3D). Thus, the most efficient means of producing allodynia for instances of the subcircuit in Cluster 1 involve disinhibition of the *E* population by lowering the response of the *I* population to *Aβ* input by reducing *ĝ*_*AβI*_, and over-exciting the *E* population by increasing *ĝ*_*AβE*_ (Fig 3E). We characterize this mechanism of disrupting the E-I balance as the *E* population being “released” from inhibitory control.

Points in Cluster 2, on the other hand, are characterized by larger *ĝ*_*AβI*_ values and smaller *ĝ*_*IE*_ and *ĝ*_*AβE*_ values (Fig 3B), relative to all points in the APS. Points in Cluster 2 are typically farther from the allodynia surface, indicating that these instances of the subcircuit may be more protected against *E* -*I* disruption (Fig 3C). The shortest paths from points in Cluster 2 to the allodynia surface involve decreasing *ĝ*_*IE*_ and increasing *ĝ*_*AβE*_ (Fig 3D) in a fixed ratio, in contrast to points in Cluster 1 (see S4 Fig).

Thus, the shortest paths to the allodynia surface from points in Cluster 2 are always in exactly the same direction, namely a ∼ 30% decrease in *ĝ*_*IE*_ and a ∼ 20% increase in *ĝ*_*AβE*_. The most efficient way to induce allodynia for points in Cluster 2 involve a lowering of the inhibition onto the excitatory population through reduction of *ĝ*_*IE*_ coupled with an over-excitation of the excitatory population through an increase in *ĝ*_*AβE*_. We characterize this mechanism for allodynia as the *E* population “escaping” inhibitory control.

The differences in these mechanisms for allodynia can be illustrated by considering how population firing-rate changes in response to A*β* input for coupling strengths sampled from the allodynia surface *S*. For example, Fig 3E (top panels) shows that a typical subcircuit instantiation in Cluster 1 with parameter values drawn from the APS (solid curves) exhibits *I* population firing rates around 50 Hz during non-painful A*β* input. However, when coupling strengths are set to the associated closest point on *S, I* population firing rates are much lower. In this way, the *E* population is “released” from inhibitory control and is able to fire.

On the other hand, for a typical subcircuit instantiation in Cluster 2 (bottom panels), the *I* population firing rate is saturated at its maximum value (80 Hz) during A*β* input for coupling strengths in the APS, as well as for coupling strengths set at their *S* values. Here, *E* population firing is promoted in the allodynia case because the response of the *E* population to the inhibitory input is altered through a decreased *g*_*IE*_ (the inhibitory firing rate remains the same as in the non-painful case). Thus, the *E* population fires because it is able to “escape” inhibitory control.

For the simple subcircuit, the APS is basically evenly split into the 2 clusters with ∼52% of points in Cluster 1 and ∼48% in Cluster 2, suggesting that the release and escape mechanisms for allodynia are equally likely to occur. While these mechanisms for allodynia are intuitively clear and perhaps unsurprising, as shown below for the static and dynamic allodynia subcircuits, the most likely allodynia mechanism may be biased towards one of these mechanisms and can be generated by different subcircuit components when the subcircuit structure is more complex.

### 2.2 Analysis of the subcircuit mediating static allodynia

The static subcircuit consists of two inhibitory populations (*I1* and *I2*), and one excitatory (*E*) population with all three populations receiving *Aβ* input (Fig 4A). We assume that the *E* population directly relays signals to projection neurons and allodynia is defined as any firing of the *E* population in response to typical non-painful A*β* input. The APS is defined in the 5-dimensional space of coupling parameters (*g*_*AβI*1_, *g*_*I*1*E*_, *g*_*AβE*_, *g*_*AβI*2_, *g*_*I*2*E*_) and the allodynia surface is a hypersurface in this space.

**Fig 4.**
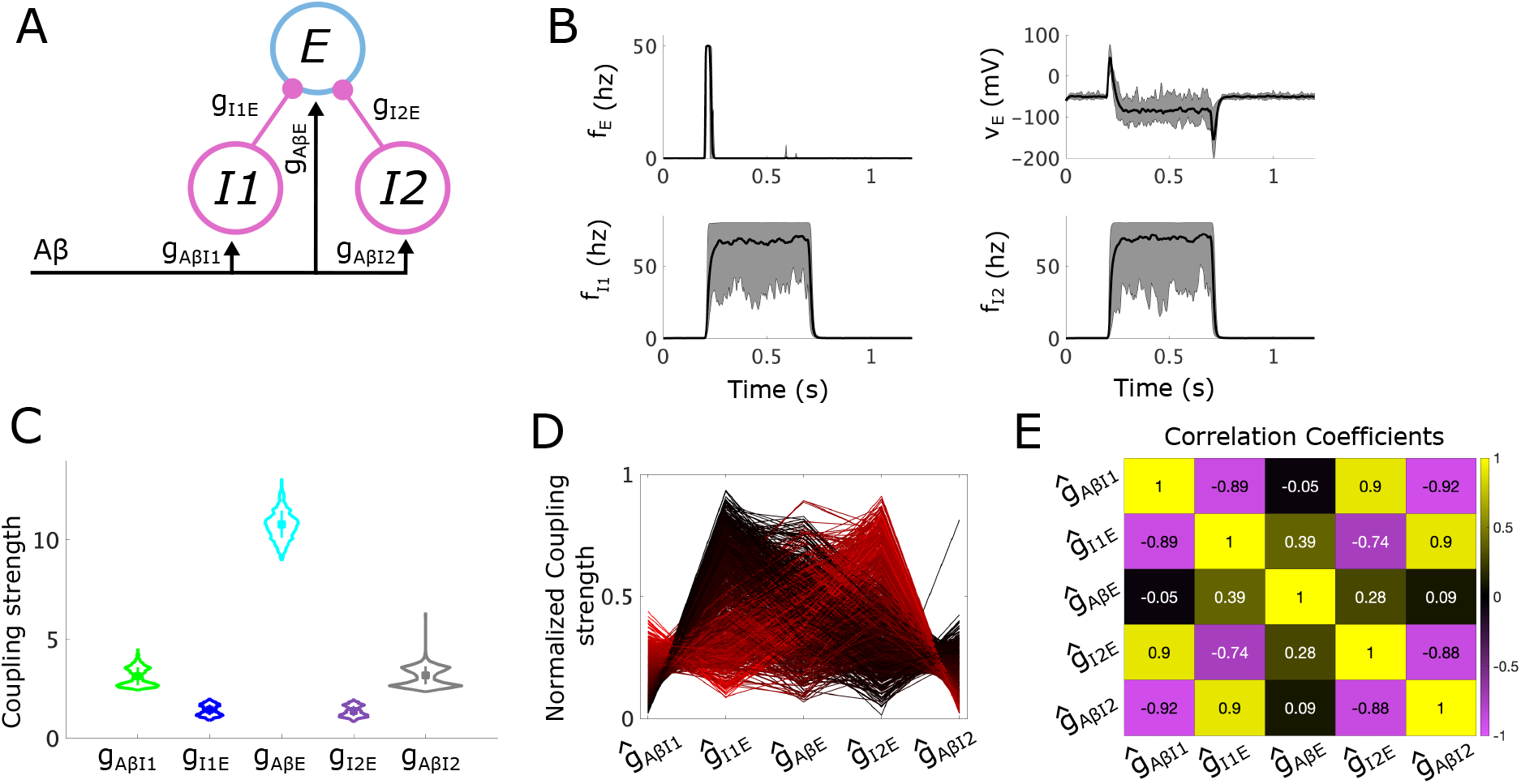
Allowable parameter space (APS) for the proposed subcircuit mediating static allodynia. **(A)** A schematic of the static subcircuit. *I1* and *I2* represent the populations of inhibitory neurons and E represents the population of excitatory neurons that synapse onto projection neurons. *Aβ* represents inputs to the subcircuit relayed from the periphery along *Aβ* fibers. **(B)** The mean (black lines) and range (shaded gray areas) for the firing rate (top left) and voltage (top right) of the *E* population, as well as for the firing rates of the *I1* (bottom left) and *I2* (bottom right) populations calculated from 20 sampled sets of coupling strengths each with a different random *Aβ* input stimulus (active during *t* ∈ [0.2, 0.7] s). **(C)** Violin plots showing the distribution of each coupling strength parameter with mean (square marker) and range of values that lie within one standard deviation (vertical bar) indicated. **(D)** Parallel plot representation of the sampled sets of normalized coupling strengths. A line gives the values of each coupling strength in a sampled set, colored on a gradient from light red to dark red according to *g*_*AβI*1_ so as to easier differentiate between individual lines. **(E)** Normalized Pearson correlation coefficients between sampled sets of normalized coupling strengths.

While the *I1* and *I2* populations represent classes of inhibitory interneurons with different molecular markers (DYN+ and PV+, respectively), we assume they have similar response properties [26] and use the same model parameters for each population. In this way, the subcircuit is symmetric in the sense that there is nothing to distinguish the two inhibitory populations from one another. As shown below, this symmetry is reflected in the structure of the APS for this subcircuit, as well as in features of the predicted most likely mechanisms for allodynia. However, in contrast to the simple subcircuit, the most likely allodynia mechanisms are biased towards different modes of *E* population release from inhibitory control.

#### The APS for the static subcircuit

To define the APS, we impose conditions on the steady-state voltages of the *E, I1*, and *I2* populations so that all instantiations of the static subcircuit display desired experimentally identified behaviors. In particular, we again require that all steady-state voltages remain within reasonable bounds. Since ablation of either inhibitory population can induce allodynia, we require both of the inhibitory populations to be active in order to maintain pain inhibition. Thus, under control conditions, typical *Aβ* stimuli induce both *I1* and *I2* firing, preventing *E* from firing, and reducing *V*_*E*_ below its resting voltage. However, if the *I1* population is ablated, then the excitatory signaling from *Aβ* input is strong enough to overcome the remaining inhibition from *I2* to induce *E* firing. Likewise, if *I2* is ablated, we expect *E* to fire. We do not incorporate ablation of both *I1* and *I2* in developing our conditions for the static subcircuit, as that would lead to extreme over-excitation of the *E* population.

Each of these conditions results in an inequality on steady-state population voltages, summarized in Table 2, that can be re-written as a set of inequalities for the coupling strengths that must be satisfied for all *f*_*Aβ*_ input levels in the range [10, 20] Hz (see Table 7). The APS for the static subcircuit is then defined as the sets of 5-tuples of coupling strengths (*g*_*AβI*1_, *g*_*I*1*E*_, *g*_*AβE*_, *g*_*AβI*2_, *g*_*I*2*E*_) which satisfy the system of inequalities and optimization problems. As discussed in Section 4.4.2, we simplify the optimization problems using Lambert functions and then uniformly sample from the defined APS using our customized sampling algorithm (Section 4.6).

**Table 2.** Conditions that the proposed subcircuit mediating static allodynia must satisfy and the resulting inequalities on steady-state voltages. We ensure that the subcircuit exhibits these behaviors by imposing conditions (middle column) on the subcircuit exhibited in either control, I1-ablation, or I2-ablation conditions (left-most column), and is realized as an inequality (right-most column) on the steady-state voltage of a population.

| Condition Type | Condition | Steady state voltage inequality—for all $f_{A\beta}$ |
| --- | --- | --- |
| Control conditions | $V_{I1}$ upper bound | $V_{I1,max} \geq V_{I1}^{ss} = g_{A\beta I1} f_{A\beta} + V_{I1,rest}$ |
| | $I1$ fires | $V_{I1,thr} \leq V_{I1}^{ss} = g_{A\beta I1} f_{A\beta} + V_{I1,rest}$ |
| | $V_{I2}$ upper bound | $V_{I2,max} \geq V_{I2}^{ss} = g_{A\beta I2} f_{A\beta} + V_{I2,rest}$ |
| | $I2$ fires | $V_{I2,thr} \leq V_{I2}^{ss} = g_{A\beta I2} f_{A\beta} + V_{I2,rest}$ |
| | Pain inhibition | $V_{E,rest} \geq V_E^{ss} = g_{A\beta E} f_{A\beta} - g_{I1E} f_{I1} - g_{I2E} f_{I2} + V_{E,rest}$ |
| | $V_E$ lower bound | $V_{E,min} \leq V_E^{ss} = g_{A\beta E} f_{A\beta} - g_{I1E} f_{I1} - g_{I2E} f_{I2} + V_{E,rest}$ |
| I1 ablation conditions | E fires | $V_{E,thr} \leq V_E^{ss} = g_{A\beta E} f_{A\beta} - g_{I2E} f_{I2} + V_{E,rest}$ |
| | $V_E$ upper bound | $V_{E,max} \geq V_E^{ss} = g_{A\beta E} f_{A\beta} - g_{I2E} f_{I2} + V_{E,rest}$ |
| I2 ablation conditions | E fires | $V_{E,thr} \leq V_E^{ss} = g_{A\beta E} f_{A\beta} - g_{I1E} f_{I1} + V_{E,rest}$ |
| | $V_E$ upper bound | $V_{E,max} \geq V_E^{ss} = g_{A\beta E} f_{A\beta} - g_{I1E} f_{I1} + V_{E,rest}$ |

We proceed with the analysis just as we did for the simple subcircuit. We first illustrate the response of subcircuit instantiations in the APS to typical non-painful *Aβ* input by simulating instantiations of the subcircuit, defined by 20 points randomly sampled from the APS (Fig 4B). Subcircuit responses show that the firing rates of the *I1* and *I2* populations always rise to at least 25% of the maximum firing rate and often approach the 80 Hz maximum. The resulting inhibition from *I1* and *I2* prevents the *E* population from firing, causing the *E* voltage to drop, as required for pain inhibition.

Violin plots (Fig 4C) of the distributions of coupling strengths in the APS highlight the symmetry of the static subcircuit. In particular, the distribution of *g*_*AβI*1_ is very similar to that of *g*_*AβI*2_. Likewise, the distribution of *g*_*I*1*E*_ is very similar to the distribution of *g*_*I*2*E*_. Also notable is that the range of *g*_*AβE*_ is largest, followed by the ranges of *g*_*AβI*1_ and *g*_*AβI*2_. The smallest ranges belong to *g*_*I*1*E*_ and *g*_*I*2*E*_, suggesting that the subcircuit is least sensitive to changes in *g*_*AβI*1_, *g*_*AβI*2_, and *g*_*AβE*_, while most sensitive to changes in *g*_*I*1*E*_ and *g*_*I*2*E*_. In addition, values of *g*_*AβE*_ in the APS are far larger than in the simple subcircuit, reflecting the need for *g*_*AβE*_ to balance out two sources of inhibition under control conditions, as well as to overcome the inhibition from one inhibitory population under ablation of either the *I1* or *I2* population.

Further, as seen in Fig 4C and Fig 4D, there are a few points with values of *g*_*AβI*1_ and *g*_*AβI*2_ much larger than is typical. This indicates that there are long and narrow regions of the APS with very low-volume, appearing for high values of *g*_*AβI*1_ and *g*_*AβI*2_. In the corresponding subcircuit instantiations, firing rate responses of *I1* and *I2* would be very strong, but because *g*_*I*1*E*_ and *g*_*I*2*E*_ are very small, the impact of inhibitory signaling on the *E* population is likely relatively weak.

We also see from the violin plots that values of *g*_*AβI*1_, *g*_*AβI*2_, *g*_*I*1*E*_, and *g*_*I*2*E*_ are strongly bimodal. This bimodality is reflected in the two trends seen in the parallel plot (Fig 6D) representing the sampled sets of coupling strengths in normalized parameter space. In one trend, *ĝ*_*I*1*E*_ is large (blacker lines), whereas the other trend (redder lines) involves larger *ĝ*_*I*2*E*_ values. From the parallel plot, we can also begin to see a number of correlations between coupling strengths. For instance, when *ĝ*_*AβI*1_ is large, *ĝ*_*I*1*E*_ is small, and similarly, when *ĝ*_*AβI*2_ is large, *ĝ*_*I*2*E*_ is small. Additionally, when *ĝ*_*I*1*E*_ is large, *ĝ*_*I*2*E*_ is small. These correlations are confirmed by the Pearson’s correlation coefficients between each normalized coupling strength (Fig 6E). Generally these correlations reflect an E-I balance manifested as the excitatory (*Aβ*) inputs to the excitatory population balanced by a combination of inputs from the two inhibitory populations. Specifically, inputs from *I1* and *I2* compensate for each other where if the inhibition from *I1* is large, then the inhibition from *I2* is small.

#### Mechanisms of allodynia in the static subcircuit

To identify sensitivities to allodynia of the static subcircuit, we consider the allodynia surface *S*_*stat*_, which in normalized parameter space is the set of points in (*ĝ*_*AβI*1_, *ĝ*_*I*1*E*_, *ĝ*_*AβE*_, *ĝ*_*AβI*2_, *ĝ*_*I*2*E*_)-space above which subcircuit instantiations produce allodynia for at least some typical *f*_*Aβ*_ (Section 4.7). We then compute the shortest vectors from each sampled point in the APS to *S*_*stat*_. Clustering of the APS based on the direction of the shortest vectors using density-based scanning (see Section 4.9) indicates that the APS divides into four clusters: Cluster 1 (green), Cluster 2 (blue), Cluster 3 (cyan), and Cluster 4 (gray) (Fig 5).

**Fig 5.**
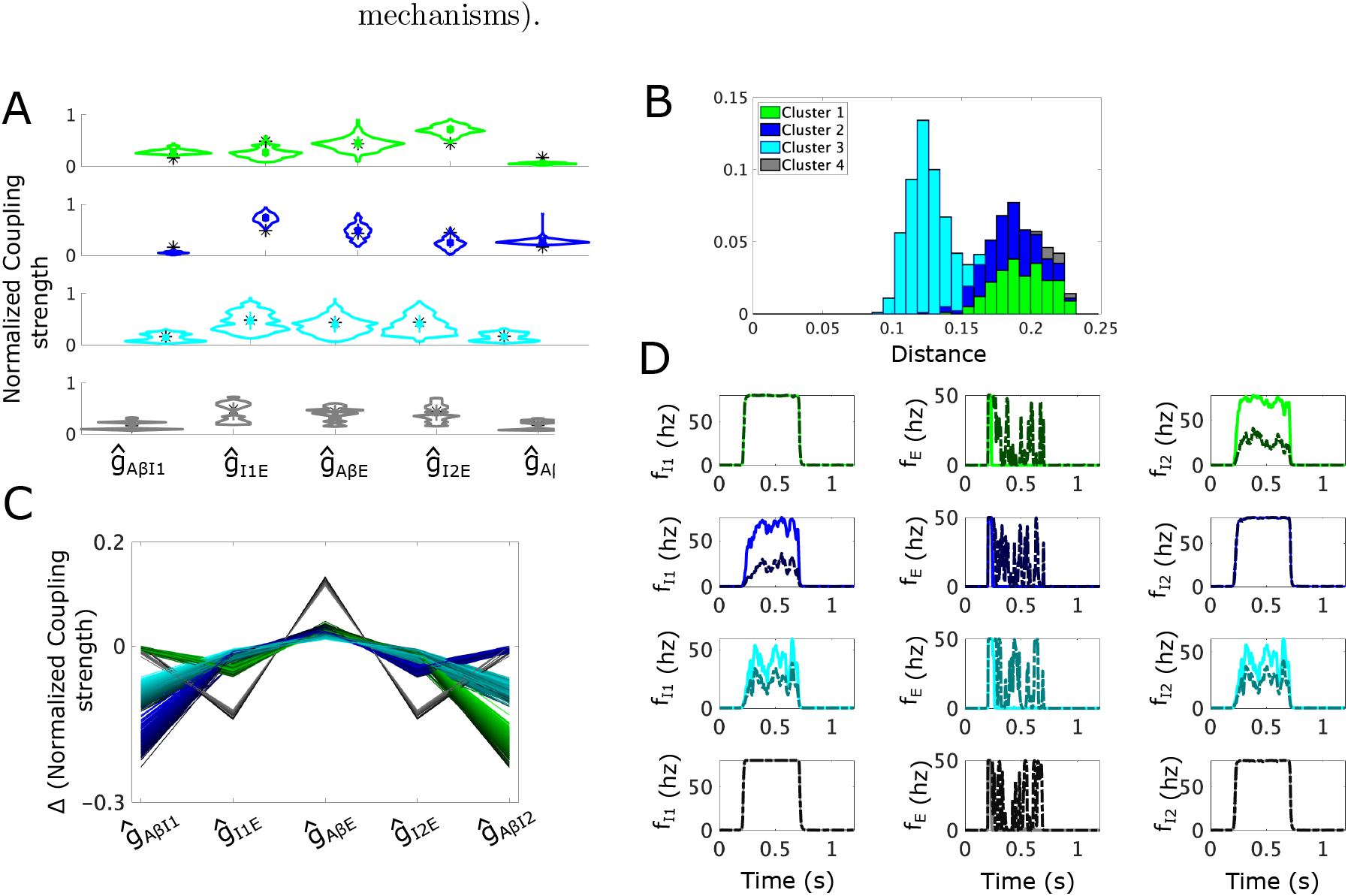
Mechanisms for static allodynia represented as shortest paths to the allodynia surface for the static allodynia subcircuit. **(A)** Violin plots for coupling strength distributions in the 4 clusters of shortest path lengths to the static allodynia surface: Cluster 1 (green, top panel), Cluster 2 (blue, 2nd panel), Cluster 3 (cyan, 3rd panel), and Cluster 4 (gray, bottom panel). Black ∗represents the mean APS coupling strength value, colored square is mean value for the cluster. **(B)** Probability distribution of the shortest distances to the allodynia surface (overall profile); shading represents the contributions from each cluster to the overall distribution. For instance, a bar that is 70% light blue indicates that cluster 3 constitutes 70% of subcircuit instantiations with the corresponding distance to the allodynia surface. **(C)** Parallel plot representation of the shortest paths to the allodynia surface from each sampled point from the APS. A line gives the components of the displacement vector corresponding to one such shortest path. Lines are colored based on clusters with a color gradient for better visualization. **(D)** Firing-rate responses to a noisy *Aβ* input signal (active during *t* ∈ [0.2, 0.7] s) for each population (columns) and for each cluster (rows). Solid lines correspond to a subcircuit instantiation with coupling strengths given by the mean values for the particular cluster (except for Cluster 4, which is spatially disjoint, where we use a representative sampled set of clustering strengths), and dash-dotted lines correspond to the subcircuit instantiations with coupling strengths given by the corresponding closest point on the allodynia surface. *Aβ* input frequencies are chosen as the smallest value that induces allodynia for each cluster.

**Fig 6.**
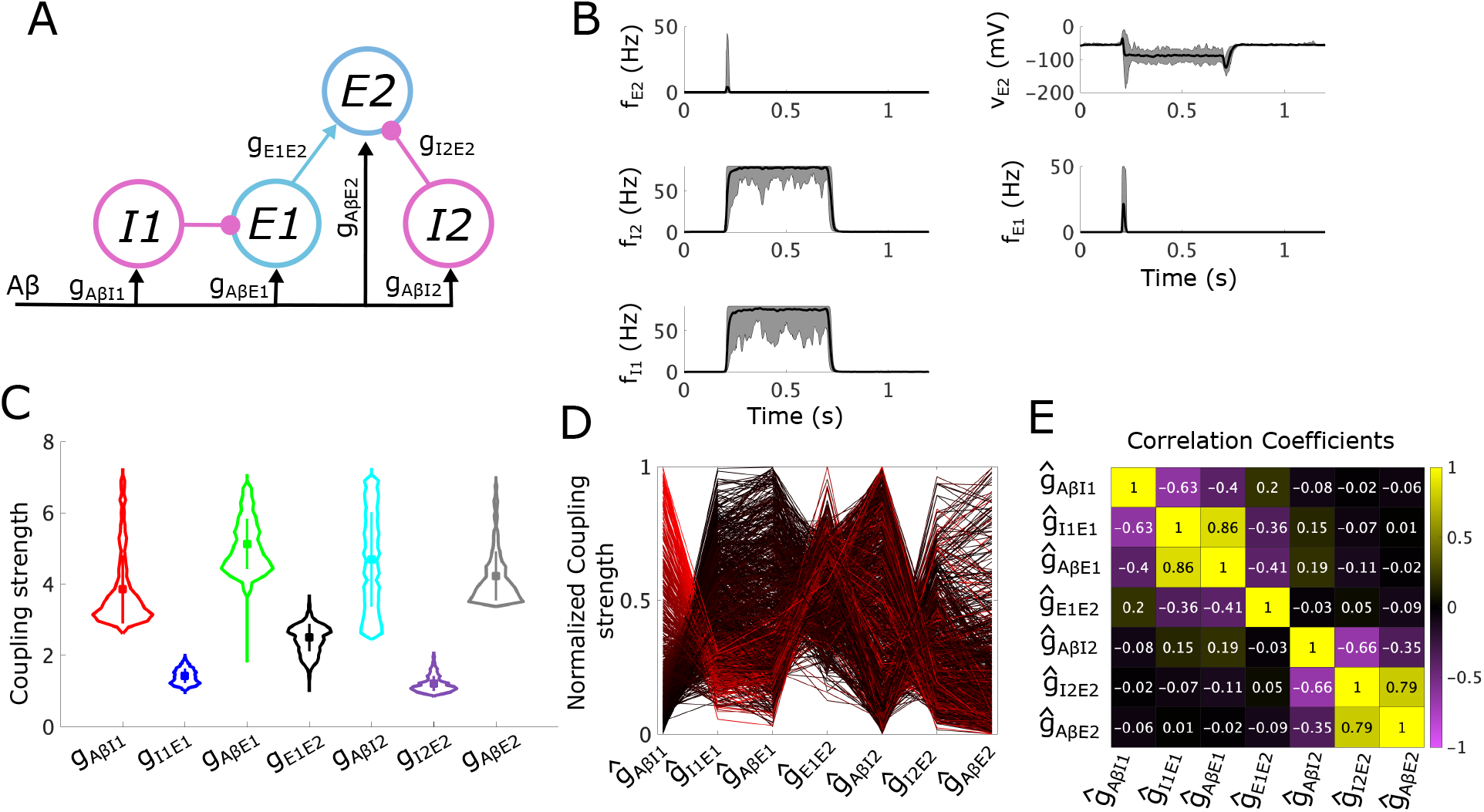
Allowable parameter space (APS) for the proposed subcircuit mediating dynamic allodynia. **(A)** A schematic of the dynamic subcircuit. *I1* and *I2* represent the populations of inhibitory neurons, and *E1* and *E2* represent the populations of excitatory neurons. We assume the *E2* population relays signals to projection neurons. *Aβ* represents inputs to the subcircuit, relayed from the periphery along *Aβ* fibers. **(B)** The mean (black lines) and range (shaded green areas) for the firing rate (top left) and voltage (top right) of the *E2* population, as well as for the firing rates of the I2 (middle left), E1 (middle right), and I1 (bottom left) populations calculated across 20 sampled sets of coupling strengths each with a different random input *Aβ* stimulus (active during *t* ∈ [0.2, 0.7] s). **(C)** Violin plots of distributions of coupling strength values in the APS. The vertical bar represents the values within one standard deviation of the mean (square marker) for the corresponding coupling strength. **(D)** Parallel plot representation of the sampled sets of normalized coupling strengths. A line gives the values of each coupling strength in a sampled set, colored on a gradient from light red to dark red according to *g*_*AβI*1_. **(E)** Normalized Pearson’s correlation coefficients between coupling strength values across sampled sets in the APS.

Cluster 1 (green) is characterized by larger *ĝ*_*AβI*1_ values and smaller *ĝ*_*I*1*E*_ values compared to the APS mean (Fig 5A, top panel). Conversely, *ĝ*_*AβI*2_ values are smaller and *ĝ*_*I*2*E*_ values are larger than the APS mean. This reflects that subcircuit instantiations in Cluster 1 display high *I1* population activity that weakly inhibits the *E* population, and a weakly active *I2* population whose inhibitory effect on *E* is strong.

For the subcircuit instantiations in Cluster 1, most efficiently reaching the allodynia surface primarily involves a decrease in *ĝ*_*AβI*2_ values (Fig 5C). This parameter change reduces the weak *I2* population activity and releases the *E* population from the inhibitory control provided by *I2*. The shortest vectors to *S*_*stat*_ simultaneously involve small decreases in *ĝ*_*I*1*E*_ values and small increases in *ĝ*_*AβE*_ values. These parameter changes additionally promote *E* population firing through an escape mechanism by slightly reducing the inhibitory effect of *I1* activity and slightly increasing *E* responses to excitatory A*β* input. The asymmetry in the inhibitory control of the *E* population under control conditions is maintained in the mechanism for allodynia in that the more weakly active *I2* population becomes weaker and the more strongly active *I1* population is not affected. This is evident in the numerical simulations of Cluster 1 subcircuit instantiations shown in Fig 5D (top row). In particular, a typical Cluster 1 subcircuit instantiation exhibits a decrease in *I2* activity in the allodynia condition compared to control, but no change in *I1* activity.

Subcircuit instantiations in Cluster 2 (blue) exhibit the opposite asymmetry in inhibitory control of the *E* population with higher *ĝ*_*AβI*2_ values compared to *ĝ*_*AβI*1_ and lower *ĝ*_*I*2*E*_ values compared to *ĝ*_*I*1*E*_ (Fig 5A, 2nd panel from the top). Thus, in these subcircuit instantiations, the *I1* population is weakly active compared to *I2* and the direction of the shortest vectors to *S*_*stat*_ primarily involves decreases in *ĝ*_*AβI*1_. This results in lower *I1* activity in the allodynia condition and release of the *E* population from *I1* inhibitory control (Fig 5C and D). Clusters 1 and 2 combined make up about half of the APS points and they display approximately the same mean distances to the allodynia surface (Fig 5B).

While subcircuit instantiations in Clusters 1 and 2 both exhibit asymmetry in the activity levels of the two inhibitory populations, in Cluster 3 (cyan) the inhibitory control of the *E* population is generally equally shared by *I1* and *I2*. Indeed, the coupling strength distributions are very similar between *ĝ*_*AβI*1_ and *ĝ*_*AβI*2_, as well as between *ĝ*_*I*1*E*_ and *ĝ*_*I*2*E*_ (Fig 5A, third panel). Cluster 3 is the largest cluster and its points lie closest to the allodynia surface (Fig 5B), suggesting that an equally distributed inhibitory gate on *E* activity is the most likely configuration under our conditions, but that it is also most easily disrupted.

To induce allodynia in Cluster 3 subcircuit instantiations, it is most efficient to decrease both *ĝ*_*AβI*1_ and *ĝ*_*AβI*2_ by roughly equal amounts and to increase *ĝ*_*AβE*_ (Fig 5C), reflecting release from both *I1* and *I2* inhibitory control. As shown in Fig 5D, (third row), for typical subcircuit instantiations in Cluster 3, neither *I1* or *I2* firing rates are at their maximum values in control or allodynia conditions, and slight decreases in both their firing rates is sufficient to allow *E* firing in the allodynia condition.

Clusters 1, 2, and 3 make up almost all of the APS, but ∼2% falls into Cluster 4 (gray). Points in Cluster 4 are also symmetric in the distribution of coupling strengths related to the inhibitory populations, and they are more bimodal than the APS as a whole. This is particularly reflected in the distributions of *ĝ*_*AβI*1_ and *ĝ*_*AβI*2_ (Fig 5A, bottom panel), suggesting that Cluster 4 has at least two groups of subcircuit instantiations that are far apart in parameter space, as is evident in the parallel plot displayed in S3 Fig. In fact, these two groups show similarities to parameter values in Cluster 1 or in Cluster 2, respectively.

However, unlike Clusters 1 and 2, allodynia is most easily induced for all Cluster 4 subcircuit instantiations by decreasing both *ĝ*_*I*1*E*_ and *ĝ*_*I*2*E*_ and increasing *ĝ*_*AβE*_, reflecting an escape of the *E* population from inhibitory control (Fig 5C). As shown in Fig 5D, (bottom panel), a typical subcircuit instantiation in Cluster 4 has inhibitory populations saturated near the maximum firing-rates, despite the asymmetry mentioned above. Thus, decreasing *ĝ*_*I*1*E*_ and *ĝ*_*I*2*E*_ reduces the effect of the two inhibitory populations. The fact that Cluster 4 is very small and its subcircuit instantiations are considerably further from the allodynia surface compared to the other clusters (Fig 5B) suggests that this mechanism for E-I balance and its disruption is less likely to occur in this subcircuit structure.

In summary, the most likely mechanisms for allodynia in the static subcircuit involve release of the *E* population from inhibitory control where the locus of decreased inhibitory signaling can occur in different parts of the subcircuit. The largest cluster (Cluster 3), and thus, the predicted most likely mechanism, is when activity levels are similar in the *I1* and *I2* populations, and their simultaneous decrease releases the *E* population to activate in response to non-painful A*β* input. The other more likely mechanisms (Cluster 1 and Cluster 2) occur when one of the inhibitory populations activates more weakly than the other in control conditions, and allodynia occurs when the activity levels of the weaker population decrease further. These results suggest that when inhibitory gating of excitatory cell activity is distributed among separate neuronal populations, the disruption of E-I balance is biased towards reduced inhibitory signaling (i.e. release mechanisms) compared to enhanced excitatory cell responses (i.e. escape mechanisms).

### 2.3 Analysis of the subcircuit mediating dynamic allodynia

In the dynamic subcircuit, allodynia is defined as firing of the *E2* population in response to typically non-painful A*β* input. In this case, the APS is defined in the 7-dimensional space of coupling strength parameters (*g*_*AβI*1_, *g*_*I*1*E*1_, *g*_*AβE*1_, *g*_*E*1*E*2_, *g*_*AβI*2_, *g*_*I*2*E*2_, *g*_*AβE*2_) and the allodynia surface is a hypersurface in this space. The structure of the dynamic subcircuit consists of two simple subcircuits coupled together, one consisting of *E1* and *I1*, and the other consisting of *E2* and *I2* (Fig 6A). However, our analysis shows that the most likely allodynia mechanisms do not exhibit the same symmetries as in the simple subcircuit; instead the escape from inhibition mechanism dominates.

#### The APS for the dynamic subcircuit

Again, to define the APS, we impose conditions on the steady-state voltages of the *E1, E2, I1*, and *I2* populations so the dynamic subcircuit displays experimentally-observed behaviors under normal healthy conditions (Table 3). Specifically, we again require that all steady-state voltages remain within reasonable bounds. In response to A*β* input in the typically non-painful range (i.e. *f*_*Aβ*_ ∈ [10, 20] Hz), the *I1* and *I2* populations should fire and provide inhibitory control of the activity of the *E1* and *E2* populations, respectively. In particular, to account for the phenomenon of pain inhibition, we require that steady-state voltages of both *E1* and *E2* are hyperpolarized from resting voltage by this inhibitory input. However, if the *E1* population is ablated, the *E2* voltage remains within the reasonable bounds. We further require that if the *I1* population is ablated, then the *E1* population fires in response to typical *Aβ* stimuli, and in turn excites the *E2* population to firing with both their voltages remaining within the reasonable bounds. Finally, if the *I2* population is ablated, then we require that the *E2* population fires in response to typical *Aβ* stimuli.

**Table 3.** Conditions that the proposed subcircuit mediating dynamic allodynia must satisfy and the resulting inequalities on steady-state voltages in response to *Aβ* input. We ensure that the subcircuit exhibits these behaviors by imposing conditions (middle column) on the subcircuit. Each condition is exhibited in either control, E1-ablation, I1-ablation, or I2-ablation conditions (left-most column), and is realized as an inequality (right-most column) on the steady-state voltage of a population.

| Condition Type | Condition | Steady state voltage inequality—for all $f_{A\beta}$ |
| --- | --- | --- |
| Control conditions | $V_{I1}$ upper bound | $V_{I1,max} \geq V_{I1}^{ss} = g_{A\beta I1} f_{A\beta} + V_{I1,rest}$ |
| | $I1$ fires | $V_{I1,thr} \leq V_{I1}^{ss} = g_{A\beta I1} f_{A\beta} + V_{I1,rest}$ |
| | $V_{I2}$ upper bound | $V_{I2,max} \geq V_{I2}^{ss} = g_{A\beta I2} f_{A\beta} + V_{I2,rest}$ |
| | $I2$ fires | $V_{I2,thr} \leq V_{I2}^{ss} = g_{A\beta I2} f_{A\beta} + V_{I2,rest}$ |
| | Pain inhibition (E2) | $V_{E2,rest} \geq V_{E2}^{ss} = g_{A\beta E2} f_{A\beta} + g_{E1 E2} f_{E1} - g_{I2 E2} f_{I2} + V_{E2,rest}$ |
| | Pain inhibition (E1) | $V_{E1,rest} \geq V_{E1}^{ss} = g_{A\beta E1} f_{A\beta} - g_{I1 E1} f_{I1} + V_{E1,rest}$ |
| E1 ablation conditions | $V_{E2}$ lower bound | $V_{E2,min} \leq V_{E2}^{ss} = g_{A\beta E2} f_{A\beta} - g_{I2 E2} f_{I2} + V_{E2,rest}$ |
| I1 ablation conditions | E1 fires | $V_{E1,thr} \leq V_{E1}^{ss} = g_{A\beta E1} f_{A\beta} + V_{E1,rest}$ |
| | $V_{E2}$ upper bound | $V_{E2,max} \geq V_{E2}^{ss} = g_{A\beta E2} f_{A\beta} + g_{E1 E2} f_{E1} - g_{I2 E2} f_{I2} + V_{E2,rest}$ |
| | E2 fires | $V_{E2,thr} \leq V_{E2}^{ss} = g_{A\beta E2} f_{A\beta} + g_{E1 E2} f_{E1} - g_{I2 E2} f_{I2} + V_{E2,rest}$ |
| I2 ablation conditions | $V_{E2}$ upper bound | $V_{E2,max} \geq V_{E2}^{ss} = g_{A\beta E2} f_{A\beta} + g_{E1 E2} f_{E1} + V_{E2,rest}$ |
| | E2 fires | $V_{E2,thr} \leq V_{E2}^{ss} = g_{A\beta E2} f_{A\beta} + g_{E1 E2} f_{E1} + V_{E2,rest}$ |

Each of these conditions leads to an inequality on steady state population voltages (Table 3) that is re-written as a set of inequalities for the coupling strengths that must be satisfied for all *f*_*Aβ*_ input levels in the range [10, 20] Hz (Table 9 in Section 4.4.3). The APS for the dynamic subcircuit is then defined as the sets of 7-tuples of coupling strengths (*g*_*AβI*1_, *g*_*I*1*E*1_, *g*_*AβE*1_, *g*_*E*1*E*2_, *g*_*AβI*2_, *g*_*I*2*E*2_, *g*_*AβE*2_) which satisfy the system of inequalities and optimization problems.

To illustrate the response of dynamic subcircuit instantiations in the APS to *Aβ* input, we simulate 20 instantiations of the subcircuit with noisy A*β* input (Fig 6B; average *f*_*Aβ*_ in [10, 20] Hz). Firing rates of the *I1* and *I2* populations rise to nearly their maximum firing rates of 80 Hz preventing *E1* and *E2* from firing and decreasing *E2* average voltage, as required for pain inhibition.

Violin plots of the sampled APS points (Fig 6C) show that the coupling strengths governing the response of the populations to A*β* input, namely *g*_*AβI*1_, *g*_*AβE*1_, *g*_*AβI*2_ and *g*_*AβE*2_, have the largest ranges, reflecting low sensitivities of these parameters for obtaining normal healthy subcircuit responses. On the other hand, the coupling strengths from the inhibitory populations to the excitatory populations, namely *g*_*I*1*E*1_ and *g*_*I*2*E*2_, have the smallest ranges, suggesting that excitatory population responses to inhibitory signaling are more constrained to maintain healthy subcircuit responses.

Correlations between coupling strength values indicate how E-I balance is maintained in the subcircuit. A parallel plot of normalized parameter sets (Fig 6D) indicates positive correlations by flat lines between two coupling strength values such as between *ĝ*_*I*1*E*1_ and *ĝ*_*AβE*1_, and between *ĝ*_*I*2*E*2_ and *ĝ*_*AβE*2_. These strong positive correlations are likewise apparent in computed correlation coefficients (Fig 6E) and reflect the balance of responses to inhibitory and excitatory inputs by the *E1* and *E2* populations, similar to what we found for the simple subcircuit. Also similar as in the simple subcircuit, negative correlations between *g*_*AβI*1_ and *g*_*I*1*E*1_, as well as between *g*_*AβI*2_ and *g*_*I*2*E*2_, indicate a preservation of inhibitory signaling to each excitatory population such that weak responses of the inhibitory populations are compensated by higher sensitivity of the excitatory populations to their activity. Further, coupling strengths pertaining specifically to the *I2* -*E2* component are only weakly correlated with coupling strengths pertaining to the *I1* -*E1* component, indicating that excitation and inhibition are being balanced separately within each subcircuit component.

#### Mechanisms of allodynia in the dynamic subcircuit

To analyze sensitivity of this subcircuit to allodynia, we consider the allodynia surface *S*_*dyn*_, which in normalized parameter space is the set of points in (*ĝ*_*AβI*1_, *ĝ*_*I*1*E*1_, *ĝ*_*AβE*1_, *ĝ*_*E*1*E*2_, *ĝ*_*AβI*2_, *ĝ*_*I*2*E*2_, *ĝ*_*AβE*2_) space above which subcircuit instantiations produce *E2* firing for at least some typical *f*_*Aβ*_ (Section 4.7). Clustering points of the APS according to the directions of the shortest vectors to *S*_*dyn*_ (Sections 4.8 and 4.9) yields four clusters (Fig 7): Cluster 1 (green), Cluster 2 (blue), Cluster 3 (red), and Cluster 4 (cyan). To summarize, Clusters 1 and 3 represent the release from inhibition allodynia mechanism in the *I2* -*E2* component and *I1* -*E1* component of the subcircuit, respectively, while Clusters 2 and 4 represent escape from inhibition in each subcircuit component, respectively.

**Fig 7.**
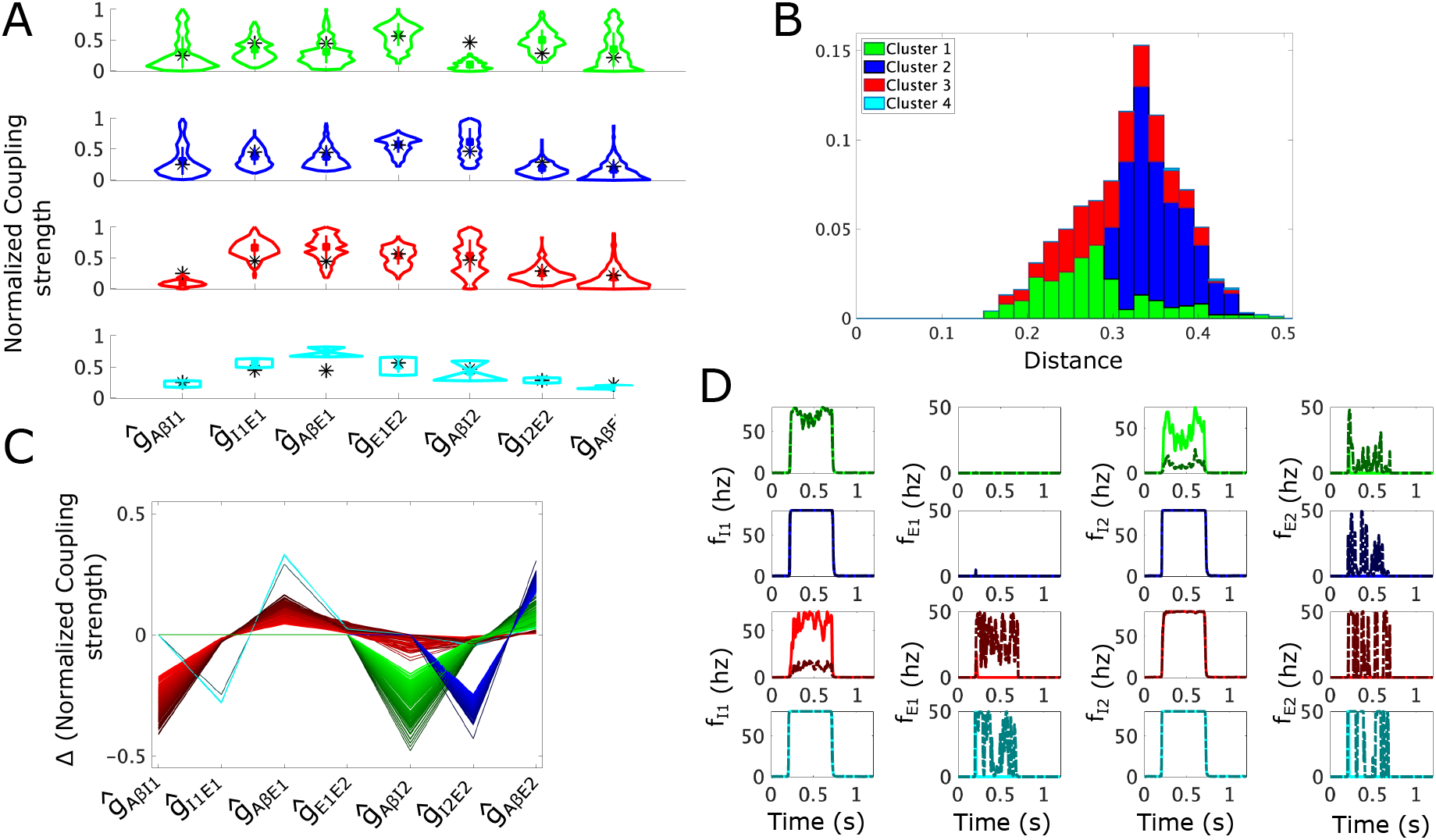
Mechanisms for dynamic allodynia represented as shortest paths to the allodynia surface for the dynamic allodynia subcircuit. **(A)** Violin plots for coupling strength distributions in the 4 clusters of shortest path lengths to the dynamic allodynia surface: Cluster 1 (green, top panel), Cluster 2 (blue, 2nd panel), Cluster 3 (red, 3rd panel), and Cluster 4 (cyan, bottom panel). Black ∗represents the mean APS coupling strength value, colored square is mean value for the cluster. **(B)** Probability distribution of the shortest distances to the dynamic allodynia surface (overall profile); shading represents the contributions of each cluster to the overall profile. For instance, a bar that is 50% dark blue indicates that cluster 2 constitutes 50% of subcircuit instantiations with the corresponding distance to the allodynia surface. Only ∼2% of sampled subcircuit instantiations belong to cluster 4 with distances greater than 3.5, thus contributing minimally to distribution. **(C)** Parallel plot representation of the shortest paths to the allodynia surface from each sampled point in the APS. A line gives the components of the shortest displacement vector from an APS point to the allodynia surface. Lines are colored based on clusters with a color gradient for better visualization.**(D)** Firing-rate responses to a noisy *Aβ* input signal (active during *t* ∈ [0.2, 0.7]) for each population (columns) and for each cluster (rows). Solid lines correspond to a subcircuit instantiation with coupling strengths given by the mean values for the particular cluster, and dash-dotted lines correspond to the subcircuit instantiation with coupling strengths given by the corresponding closest point on the allodynia surface. *Aβ* input frequencies are chosen as the smallest value that induces allodynia for each cluster.

Parameter sets belonging to Cluster 1 (green) are characterized by relatively lower *ĝ*_*AβI*2_ values and relatively higher *ĝ*_*I*2*E*2_ values compared to the APS mean values (Fig 7A), similar to Cluster 1 in the simple subcircuit. An additional similarity is that points in Cluster 1 are generally closer to *S*_*dyn*_ compared to the other clusters (Fig 7B). The shortest vectors to *S*_*dyn*_ for Cluster 1 involve traveling only in directions associated with parameters of the *I2* -*E2* component of the subcircuit (Fig 7C). Specifically, to induce allodynia primarily involves decreases in *ĝ*_*AβI*2_ leading to a dramatic reduction in the activity of *I2* in response to normal *Aβ* input and the release of the inhibitory gate on *E2*. Simulations of Cluster 1 subcircuit instantiations with parameter values on *S*_*dyn*_ show this effect (Fig 7D, top row, dash-dotted curves) where, in response to normal *Aβ* input, activity of the *I2* population is sufficiently decreased so that *E2* fires, thereby producing allodynia.

The majority of points in the APS belong to Cluster 2 (blue, Fig 7B) and, as a result, the coupling strength values for points in Cluster 2 do not greatly differ from the mean or from the overall distributions of the APS values (Fig 7A). The direction of the shortest paths to *S*_*dyn*_ involves decreasing *ĝ*_*I*2*E*2_ and increasing *ĝ*_*AβE*2_. This corresponds to an escape of *E*2 from its inhibitory control, similar to the allodynia mechanism for Cluster 2 in the simple subcircuit. This is reflected in simulations of Cluster 2 subcircuit instantiations (Fig 7D) that show that on the allodynia surface (dash-dotted curves), the *I2* population remains at highest activity levels in response to A*β* input in the normal range, but the *E2* population is able to fire.

Parameter sets in Cluster 3 differ from the mean APS values in coupling strengths associated with the *I1* -*E1* component of the subcircuit. Specifically, *ĝ*_*AβI*1_ values are lower than mean values and *ĝ*_*I*1*E*1_ and *ĝ*_*AβE*1_ values are higher (Fig 7A, red). Cluster 3 points are generally quite close to the allodynia surface, almost as close as points from Cluster 1 (Fig 7B). The direction of the shortest path to the allodynia surface involves changes in all coupling parameters but the largest parameter changes reflect an allodynia mechanism involving the release of *E*1 from inhibitory control. In particular, the largest parameter change is a decrease in *ĝ*_*AβI*1_ while increased *ĝ*_*AβE*1_, showing the next largest change, contributes to the *E*1 release mechanism (Fig 7C). Variations of the other coupling parameters in the shortest vector to *S*_*dyn*_ suggest that escape of *E*2 from its inhibitory control also contributes to this most likely mechanism for allodynia, namely a slight increase in *ĝ*_*E*1*E*2_ and slight decrease in *ĝ*_*I*2*E*2_. Simulations of Cluster 3 subcircuits with coupling strength values on *S*_*dyn*_ (Fig 7D, 3rd row, dash-dotted curves) show decreased *I1* activity that allows *E1* firing, and consequently *E2* firing as well in response to typical *Aβ* input.

Cluster 4 (cyan) represents the smallest portion of the APS (∼2%) and its points are significantly farther from the allodynia surface than the remainder of the APS. Its parameter sets are characterized by having considerably larger values of *ĝ*_*I*1*E*1_ and *ĝ*_*AβE*1_ compared to mean APS values (Fig 7A). The most efficient allodynia mechanism for Cluster 4 mainly involves escape of *E*1 from inhibitory control, but the mean shortest vector to *S*_*dyn*_ involves variation in all coupling parameters (Fig 7C). Indeed, the most allodynia-vulnerable direction for points in Cluster 4 primarily involves decreasing *ĝ*_*I*1*E*1_ and increasing *ĝ*_*AβE*1_. The other parameter variations suggest that escape of *E*2 from inhibitory control also contributes to the allodynia mechanism. Simulations of Cluster 4 subcircuits (Fig 7D, bottom row, dash-dotted curves) with parameter values on the allodynia surface show that the *E1* population fires in response to normal *Aβ* input, leading to *E2* population firing as well, while neither *I1* or *I2* show any reduction in activity.

Interestingly, while the structure of the dynamic allodynia subcircuit shows symmetry relative to the simple subcircuit, this symmetry is broken in the most likely mechanisms for allodynia. Specifically, in the simple subcircuit the escape and release from inhibition mechanisms were basically equally likely to occur across the APS. In contrast, for the dynamic subcircuit, almost half of the APS has a most likely mechanism of the *E*2 population escaping from inhibition (Cluster 2, Fig 7B). The remaining half of the APS has mechanisms split between *E*2 release from inhibition (Cluster 1) and *E*1 release from inhibition (Cluster 3), and a small portion of the APS has a mechanism of *E*1 escape from inhibition (Cluster 4). For dynamic allodynia, these results suggest that while the presence of *E*1 in the subcircuit may act as an amplifier of A*β* signaling, it may not be the most likely culprit in tipping E-I balance towards excitation to cause allodynia.

## 3 Discussion

While the basic tenets of Melzack and Wall’s “gate control” theory [10] are still pertinent, updated conceptual models for spinal cord pain signaling are needed to account for recent results identifying diverse types of excitatory and inhibitory interneurons and their circuit structure in the dorsal horn. In this study, we analyzed biophysically-motivated subcircuit models that represent common motifs in dorsal horn layer I-II pain processing neural circuits to identify the diversity of mechanisms that maintain and disrupt E-I balance in subcircuit responses to *Aβ* inputs. In our model subcircuits in normal healthy conditions, E-I balance was tuned to suppress activity of excitatory interneurons, that are presumed to directly target layer I projection neurons, in response to non-painful *Aβ* inputs. Computation of the APS for each subcircuit defined all possible models that exhibit this healthy E-I balance and also replicate experimentally observed responses to neural ablation or silencing manipulations. To identify most likely mechanisms that disrupt E-I balance in individual subcircuits, we defined allodynia surfaces and computed shortest vectors to it from each point in the APS. The direction in parameter space of the shortest vector identified the minimal alterations to the subcircuit that resulted in a disruption of E-I balance leading to activation of the excitatory interneurons driving the pain projection neurons.

Our results show that in each subcircuit, E-I balance can be disrupted by the excitatory interneurons escaping their inhibitory control or by an attenuation of inhibitory signaling that releases them from inhibitory control, also referred to as disinhibition. Many reported changes in spinal circuits induced in physiological models of chronic pain and allodynia are examples of these mechanisms. In particular, in the escape mechanism, inhibitory signaling was unaffected but its effect on post-synaptic excitatory interneurons was diminished. Physiologically, this could occur through changes in intracellular chloride concentration in post-synaptic excitatory cells that attenuates GABA-receptor mediated inhibitory synaptic currents [29, 30], down-regulation of GABA receptors or loss of synapses from inhibitory to excitatory populations [17]. Additionally, escape from inhibition could occur due to increased excitability of excitatory cells [15] or increased impact of signaling from *Aβ* fibers potentially arising from sprouting of synapses [31] or upregulation of neurotransmitter release from *Aβ* synapses [15]. In the release from inhibition mechanism, inhibitory signaling is diminished which could occur physiologically through reduced pre-synaptic GABA or glycine levels, reduced GABA or glycine release, or lower firing rates in inhibitory interneurons [15, 17, 32].

In the simple subcircuit, model analysis predicts that E-I balance dysregulation is equally likely to occur through escape or release from inhibition, but slightly higher magnitude changes are necessary for the escape mechanism. While this result is intuitively clear due to the very simple subcircuit structure, it validates that our methodology does not have any inherent biases towards either mechanism.

In the static allodynia subcircuit, inhibitory gating of the excitatory population is distributed across two distinct interneuron populations. With this structure, our results predict that the disruption of E-I balance is biased towards reduced inhibitory signaling (i.e. release mechanisms) where the locus of decreased inhibitory signaling can occur in different parts of the subcircuit. The predicted most likely mechanism, represented by the largest cluster (Cluster 3), is when both inhibitory populations contribute equally to inhibitory control of the excitatory population and a decrease in firing or signaling occurs in both populations. The next most likely mechanisms, represented by Cluster 1 and Cluster 2, occur when one of the inhibitory populations is more weakly active than the other in healthy conditions, and loss of inhibitory control occurs when the activity of the weaker population decreases further.

In the dynamic allodynia subcircuit, *Aβ* input is amplified by two distinct excitatory interneuron populations, where the *E2* population is downstream from the *E1* population. With this structure, model results predict that disruption of E-I balance is equally likely to occur through escape (Cluster 2) or release (Cluster 1 and 3) mechanisms. The site of the escape mechanism is the *E2* population, suggesting that increased activity in more downstream excitatory interneurons is more disruptive compared to similar changes in upstream excitatory interneurons. For the release mechanism, the site for loss of inhibitory control is not biased to either the upstream or downstream excitatory population but is equally likely to occur at either population.

While our analysis has centered around the minimal coupling strength changes that result in allodynia, it is important to remember that allodynia can be induced by other, higher magnitude disruptions in the subcircuit. For instance, in all three subcircuits, it is possible to induce an allodynia response by sufficiently increasing the impact of *Aβ* signaling on the most downstream excitatory population alone, so that it overcomes inhibitory control and fires in response to innocuous stimuli. Additionally, allodynia can be induced by sufficiently decreasing the impact of inhibitory signaling on the most downstream excitatory population. However, since larger magnitude changes to the subcircuit would be required in these scenarios, it may be presumed that they are less likely to occur than the minimal changes identified in our results.

### Application to more complex dorsal horn circuits

These results can be applied to help understand how E-I balance is maintained and likely disrupted in more complex dorsal horn circuits [33, 34]. In models of more complex circuits, the space of unconstrained parameters is so high-dimensional that identifying the diversity of parameter sets that satisfy desired model behaviors is difficult, even when computational optimization algorithms are implemented. Our results can provide constraints on regions of parameter space where E-I balance in subcircuits of the network is achieved. For example, the dorsal horn layer I-III neural circuit modeled by Medlock et al. [33] consists of 5 excitatory interneuron populations and two inhibitory interneuron populations with each population consisting of a network of Hodgkin-Huxley-type model neurons. In the circuit, 3 of the excitatory populations act as upstream amplifiers of *Aβ* input to 2 downstream excitatory populations that make direct connections to projection neurons. The synaptic pathways of *Aβ* input through the upstream excitatory populations to one of the downstream excitatory populations are similar to the dynamic subcircuit modeled here. Our results predict that there should be parameter sets that limit responses to *Aβ* input in the downstream excitatory population through these synaptic pathways by mechanisms reflected by the clusters found for the dynamic subcircuit. Namely, there should be parameter sets in which the downstream excitatory neuron responses are primarily gated by direct inhibition from the inhibitory cells targeting them (corresponding to Clusters 1 and 2), and parameter sets in which their responses are limited by inhibitory control of the upstream excitatory populations (corresponding to Clusters 3 and 4).

Based on Medlock et al.’s [33] finding that, in order to maintain healthy E-I balance, small reductions in inhibitory control of upstream excitatory populations required larger increases in inhibitory input to the downstream excitatory population suggests that perhaps their model parameter set may be analogous to Cluster 3 or 4 parameter sets in the dynamic subcircuit. However, since the Medlock et al. [33] network contains additional components, there may be additional constraints on achieving E-I balance that are not contained in the smaller dynamic subcircuit.

### Model limitations

A limitation of the firing rate model formalism we implemented, which only models average population voltages and firing-rates, is that specific excitability characteristics or response features that are evident on the single neuron and spike levels are not considered. Nevertheless, we did constrain the dependence of population average firing rates on membrane voltages by fitting the steady state firing rate activation functions to frequency-voltage relationships measured in dorsal horn neurons [26]. However, these relationships do not take into account some spiking patterns observed in dorsal horn excitatory interneurons, such as delayed onset of firing or transient firing [16, 20, 35, 36]. Recent development of next-generation firing rate models that can be directly reduced from networks of individual neuron models [37, 38] provide the framework to include specific spiking patterns into a mean-field reduction, such as spike frequency adaptation [39, 40]. We expect that delayed firing could similarly be accounted for in a firing-rate model reduction of simplified neuron models that are fit to observed firing patterns of dorsal horn cells. Including such spiking properties in a firing rate population network would allow analysis of the interactions of these cellular firing patterns with network structure in maintaining or disrupting E-I balance in dorsal horn circuits.

Our models and analysis focused on the response to stimuli arriving on *Aβ* fibers only and did not include activity of layer I projection neurons. The model subcircuits could be extended to explicitly include the effects of painful signals arriving on *C* fibers that directly connect to projection neurons and interneuron populations. In previous work modeling spinal subcircuits using a firing rate model formalism, we included *C* fiber input that was mediated through NMDA receptors in post-synaptic populations [41]. The NMDA-receptor mediated connection strength depended on post-synaptic average voltage to account for voltage-dependent removal of a Mg^2+^ block. Such an extended model would be able to account for wind-up of projection neuron activity in response to repetitive brief *C* fiber input and also explicitly account for pain inhibition, namely the attenuation of *C* fiber response in the projection neurons in the presence of simultaneous *Aβ* input. We note, however, that our current results would not be qualitatively affected as *Aβ* and *C* fibers are parallel pathways and their inputs do not interact.

### Application of methodology

In this study, we introduced an analysis methodology for neural population model circuits that can determine parameter values optimized to account for experimental observations and for identifying sensitivities of model behaviors to parameter variations. The methodology involves the following steps:

1. Translate normal and pathological experimental behaviors that a circuit should replicate into analytical conditions that model variables must satisfy.
2. Re-frame these conditions into systems of inequalities and optimization problems that the parameters of interest must satisfy. In our work, we focused on the parameters governing the coupling strengths between populations. These analytically determined conditions described distinct regions of parameter space in which the corresponding subcircuit instantiations displayed normal behaviors (the APS) and above which they displayed pathological behavior (above the allodynia surface).
3. Determine the most likely mechanisms that induce the pathological condition, in our case allodynia, by finding the shortest path from each parameter set in the APS to the surface at the boundary of the pathological region

This methodology can be applied to study any circuit of neural populations, although the implementation of Step 2 is more tractable for feedforward circuits with a relatively small number of populations. Thus, we expect that the methodology may be useful for analyzing propagation of signaling and E-I balance in other sensory processing circuits with feedforward structure.

### Conclusions

In all, successful treatment of chronic pain conditions, such as allodynia, require full understanding of the underlying physiological causes. Dysregulation of E-I balance in dorsal horn circuitry is a compelling cause supported by an array of pre-clinical studies, but our incomplete understanding of the circuitry and the building evidence for its complexity leave many questions for how the dysregulation occurs. The identification of multiple types of excitatory and inhibitory interneurons in dorsal horn circuits suggests that dysregulation may occur through multiple mechanisms that may be dependent on the nature of the injury or insult that induces the chronic pain condition [42]. Our modeling results identifying diverse mechanisms underlying allodynia in dorsal horn subcircuits are a first step to systematically unravel the interactions among multiple interneuron populations for the maintenance of E-I balance in healthy conditions and its disruption in allodynia. Continued experimental work identifying dorsal horn circuit structure and the functional relationships among diverse interneuron types will provide constraints necessary to construct and analyze more complete circuit models that can participate in the development of therapies for this debilitating condition.

## 4 Materials and methods

This section is organized as follows: Sections 4.1 - 4.3 contain descriptions of the subcircuit models and their parameter values; Section 4.4 defines the allowable parameter spaces for each subcircuit; Section 4.5 describes the algorithm used to normalize the APS; Section 4.6 describes the algorithm to uniformly sample the APS; mathematical descriptions of the allodynia surfaces for each subcircuit are contained in Section 4.7; and mathematical details for the computation of distance between APS points and the allodynia surface, and clustering of APS points based on that distance are in Sections 4.8 and 4.9, respectively.

### 4.1 Population firing rate model

For our models of layer I-II dorsal horn neuronal subcircuits, we implement a well-established firing rate model formalism that models the average membrane voltage and average firing rates of neuronal populations (see e.g. [23–25]). Average firing rates *f*_*x*_ are computed from average voltages *V*_*x*_ with a sigmoidal activation function of the form:

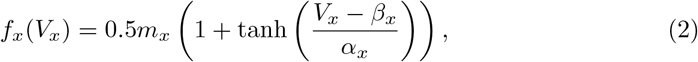

where *m*_*x*_ is the maximum firing rate of the population, *β*_*x*_ is the half-activation voltage and *α*_*x*_ governs the slope of the population’s firing-rate response to voltage changes. We match these parameters to experimental measurements of frequency-voltage relationships for dorsal horn neurons as described in Section 4.2. We expect that in the absence of inputs, *V*_*x*_ remains at a rest value *V*_*x,rest*_. In response to synaptic inputs, *V*_*x*_ deviates from its rest value according to the following differential equation:

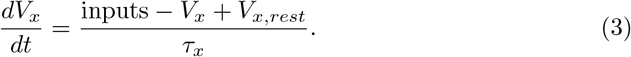

where the time constant *τ*_*x*_ describes how quickly *V*_*x*_ changes in response to inputs. The inputs are computed as the sum of the firing rates of all pre-synaptic populations to population *x*, denoted here as *y*_1_, *y*_2_, …, together with the firing rate of the *Aβ* fiber input, *f*_*Aβ*_, weighted by the corresponding coupling strengths:

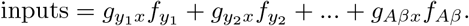

We thus expect that in the presence of steady inputs, *V*_*x*_ approaches the steady-state value given by

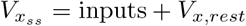

Table 4 summarizes the model equations for each subcircuit we analyze.

**Table 4.** Equations for each subcircuit model.

| Subcircuit | Population | Governing Equation |
| --- | --- | --- |
| simple subcircuit | I | $\frac{dV_I}{dt} = \frac{(g_{A\beta I}f_{A\beta} + V_{I,rest}) - V_I}{\tau_I}$ |
| | E | $\frac{dV_E}{dt} = \frac{(g_{A\beta E}f_{A\beta} - g_{IE}f_I + V_{E,rest}) - V_E}{\tau_E}$ |
| static subcircuit | I1 | $\frac{dV_{I1}}{dt} = \frac{(g_{A\beta I1}f_{A\beta} + V_{I,rest}) - V_{I1}}{\tau_I}$ |
| | I2 | $\frac{dV_{I2}}{dt} = \frac{(g_{A\beta I2}f_{A\beta} + V_{I,rest}) - V_{I2}}{\tau_I}$ |
| | E | $\frac{dV_E}{dt} = \frac{(g_{A\beta E}f_{A\beta} - g_{I1E}f_{I1} - g_{I2E}f_{I2} + V_{E,rest}) - V_E}{\tau_E}$ |
| dynamic subcircuit | I1 | $\frac{dV_{I1}}{dt} = \frac{(g_{A\beta I1}f_{A\beta} + V_{I,rest}) - V_{I1}}{\tau_I}$ |
| | E1 | $\frac{dV_{E1}}{dt} = \frac{(g_{A\beta E1}f_{A\beta} - g_{I1E1}f_{I1} + V_{E,rest}) - V_{E1}}{\tau_E}$ |
| | I2 | $\frac{dV_{I2}}{dt} = \frac{(g_{A\beta I2}f_{A\beta} + V_{I,rest}) - V_{I2}}{\tau_I}$ |
| | E2 | $\frac{dV_{E2}}{dt} = \frac{(g_{A\beta E2}f_{A\beta} + g_{E1E2}f_{E1} - g_{I2E2}f_{I2} + V_{E,rest}) - V_{E2}}{\tau_E}$ |

### 4.2 Parameters of population firing-rate models

We choose parameters for the activation functions of excitatory and inhibitory populations based on the experimental measurements of membrane properties and firing behavior in rat dorsal horn neurons reported in Ruscheweyh et al. [26]. In our subcircuits, we assume all excitatory populations have the same parameters and assume the same for inhibitory populations. We assume that average resting voltages *V*_*I,rest*_ and *V*_*E,rest*_ are ™ 60 mV, approximately the values reported in [26] Maximum firing rates of excitatory and inhibitory populations are set to 50 and 80 Hz respectively, based on [33].

We use the frequency-current relations and current-voltage relations reported in [26] to extract frequency-voltage relations. The firing-rate activation functions given in Eq (2) are fit to these frequency-voltage relations using the trust-region-reflective non-linear-least-squares algorithm via Matlab’s “fit” function [43] to obtain the values of *β*_*x*_ and *α*_*x*_ (*x* = *E, I*) for excitatory and inhibitory populations.

Maximum and minimum voltages are set to 12*α*_*x*_ mV above and below, respectively, the half-activation voltage value *β*_*x*_. The voltage thresholds for firing are defined as *β*_*x*_™*α*_*x*_. This ensures that both populations have similar behavior in response to proportional changes in input.

Finally, we choose membrane time constants so they are roughly on the same time-scale as those of [44]. Notably, since many of the results depend on steady-state voltages, small changes in the particular values of the time constants *τ*_*E*_ and *τ*_*I*_ have little effect on the qualitative behavior of the results. We choose *τ*_*E*_ = 0.01 seconds and *τ*_*I*_ = 0.02 seconds. All model parameters are listed in Table 5.

**Table 5.**
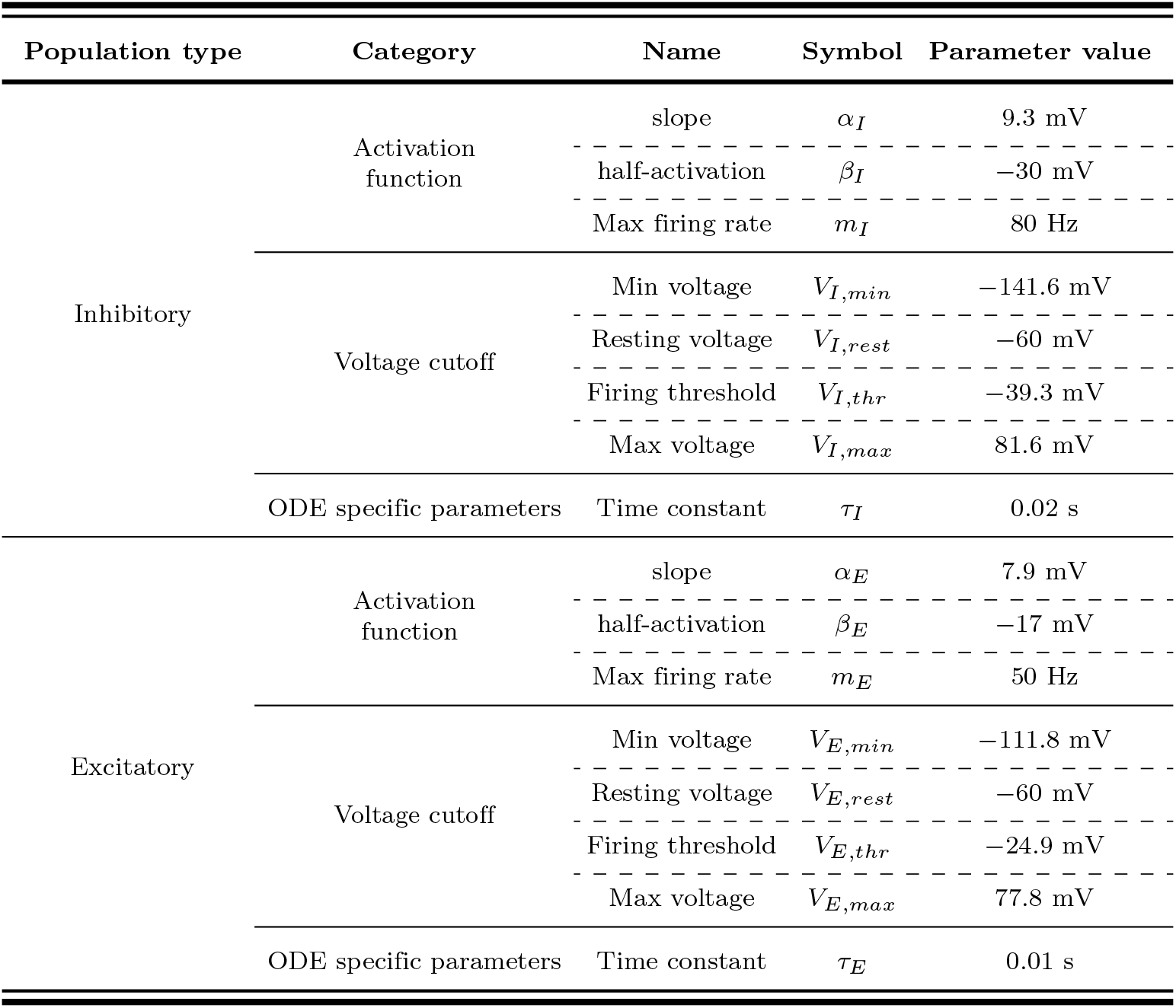
Model parameter values for inhibitory and excitatory populations. Parameters for each population type (first column) are divided into 3 categories (2^nd^ column): those that pertain to activation functions, those that give voltage cutoffs, and those that appear in the model differential equations. All these parameter values are fixed for each inhibitory or excitatory population.

### 4.3 *Aβ* stimuli model

In simulations of subcircuit responses to time-varying input on A*β* fibers, we drive neural populations with input representing the average firing rate of a bundle of 300 *Aβ* fibers (which is on the order of magnitude of the number of *Aβ* fibers in afferent nerve fibers from rat skeletal muscle, as in e.g [45]). In particular, we describe the spiking activity on each *Aβ*-fiber in the bundle via a Poisson process with a variable rate depending on the presence or absence of peripheral stimuli. Namely, in the absence of stimuli, each fiber transmits action potentials at a background rate of 1 Hz, while in the presence of a stimulus the rate increases to *f*_*Aβ*_ Hz. The input to subcircuit populations is the average spiking rate across all A*β* fibers (in Hz), namely a noisy time-varying input with average firing rate *f*_*Aβ*_ Hz. In all subcircuit simulations, stimuli are applied from 0.2 ™ 0.7 seconds.

In the parameter sensitivity analysis of model subcircuits, *f*_*Aβ*_ is taken as a constant value between [10, 20] Hz. This range of A*β* fiber firing activity has been observed in slowly adapting mechanoreceptors in response to ramp-and-hold mechanical stimulation [27].

### 4.4 Defining the allowable parameter space (APS) for the subcircuits

Our parameter sensitivity analysis method consists of translating normal experimental behaviors that the subcircuits should replicate into analytical conditions on model variables, specifically on average voltages. These conditions are then re-written into systems of inequalities and optimization problems that coupling strength parameters must satisfy. The parameter sets that satisfy these systems constitute the allowable parameter space (APS). In this section, we derive these systems for our model subcircuits. Full details of the derivation are shown for the simple subcircuit only, as the derivation follows similarly for the static and dynamic subcircuits.

#### 4.4.1 Simple subcircuit

For the simple subcircuit, it is possible to make considerable progress towards deriving an explicit definition of the allowable parameter space. As described in Table 1 in Section 2.1, the conditions on average voltages that ensure the subcircuit replicates experimentally appropriate behaviors are given as follows:

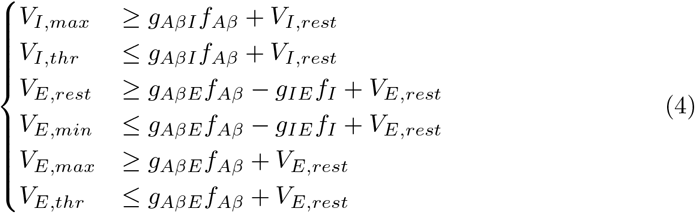

The first two inequalities can be rewritten to yield bounds on *g*_*AβI*_ :

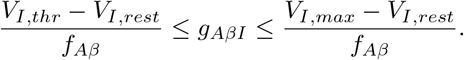

However, as these conditions must hold for all *f*_*Aβ*_ ∈ [10, 20] Hz, we need that

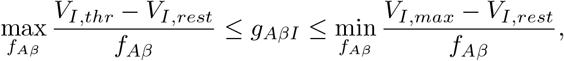

which can be re-written as

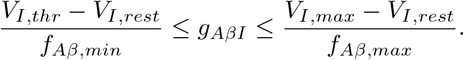

Likewise, the last two inequalities in Eq (4) yield analogous bounds on *g*_*AβE*_ and the middle two inequalities yield analogous bounds also on *g*_*AβE*_. We summarize the resulting system of inequalities for coupling strength parameters in Table 6.

**Table 6.** Inequalities on coupling strengths that define the allowable parameter space for the simple subcircuit. Inequalities on simple subcircuit coupling strength parameters that define the subcircuit APS and are obtained from the inequalities on population voltages in Table 1. (UB = upper bound, LB = lower bound, abl = ablation, PI = pain inhibition)

| Population | Condition | Inequalities |
| --- | --- | --- |
| I | UB + fires | $\frac{V_{I,thr}-V_{I,rest}}{f_{A\beta,min}} \leq g_{A\beta I} \leq \frac{V_{I,max}-V_{I,rest}}{f_{A\beta,max}}$ |
| E | UB + fires (I1 abl.) | $\frac{V_{E,thr}-V_{E,rest}}{f_{A\beta,min}} \leq g_{A\beta E} \leq \frac{V_{E,max}-V_{E,rest}}{f_{A\beta,max}}$ |
| | LB + PI | $\max_{f_{A\beta}} \frac{g_{IE}f_I - (V_{E,rest}-V_{E,min})}{f_{A\beta}} \leq g_{A\beta E} \leq \min_{f_{A\beta}} \frac{g_{IE}f_I}{f_{A\beta}}$ |

**Table 7.**
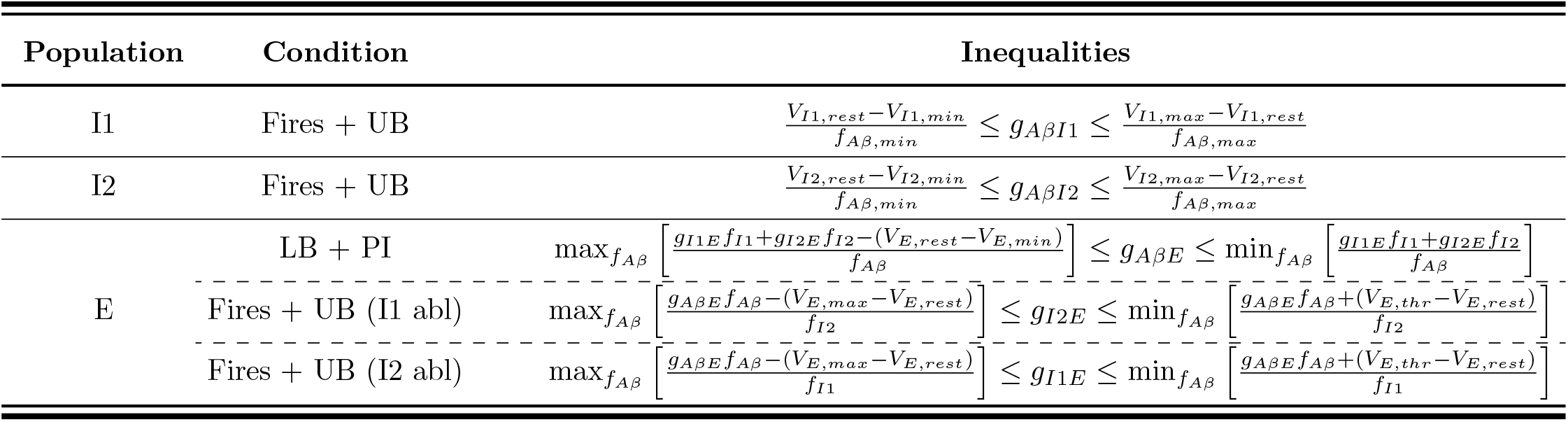
Inequalities expressed as upper and lower bounds on coupling strengths define the allowable parameter space for the static subcircuit. These inequalities are obtained by algebraically manipulating the inequalities on the voltages of various populations from Table 2 that define the APS for the static subcircuit so that the inequalities are written explicitly in terms of coupling strengths. (UB = upper bound, LB = lower bound, abl = ablation, PI = pain inhibition).

Since *f*_*I*_ is a nonlinear (hyperbolic tangent) function of *f*_*Aβ*_ and *g*_*AβI*_, the optimization problems in the last line of Table 6 are generally difficult to solve explicitly. Nevertheless, we can simplify the maximization problem – 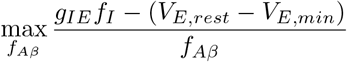 – and in fact explicitly solve the minimzation problem – 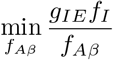 – making it much easier to numerically approximate the APS. To do so, note that both of those optimization problems can be rewritten as:

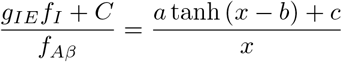

where

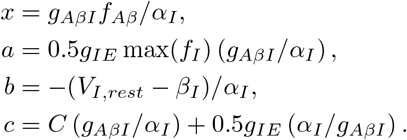

Consequently, the solutions to the preceding optimization problems occur either at *x* = *g*_*AβI*_*f*_*Aβ,min*_*/α*_*I*_, *g*_*AβI*_*f*_*Aβ,max*_*/α*_*I*_, or at one of the critical points given by

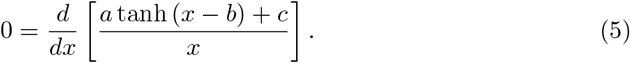

We show in S1.1 Appendix that when *C* = 0, the solutions of Eq 5 are given in terms of the 0^th^, *W*_0_, and ™1^st^, *W*_™1_, branches of the Lambert-W function:

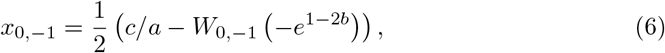

which yields

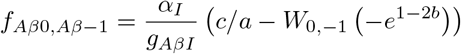

so long as *f*_*Aβ*0_ or *f*_*Aβ*™1_ ∈ [*f*_*Aβ,min*_, *f*_*Aβ,max*_] or at *f*_*Aβ*_ = *f*_*Aβ,min*_ or *f*_*Aβ*_ = *f*_*Aβ,max*_.

In S1.1 Appendix,we address the case when *C* ≠ 0 by showing how to take advantage of the structure of the optimization problem to solve it numerically.

As an alternative approach, it is possible to address the case where *C* ≠ 0 by rewriting the middle two equations of Eq 4 so they bound *g*_*IE*_ (see S1.2 Appendix). The resulting optimization problems may then be solved explicitly in terms of the Lambert W functions. While we do not implement this alternative approach for the simple circuit, we do apply it in our analysis of the static subcircuit (see below).

#### 4.4.2 Static subcircuit

For the static subcircuit, the inequalities that population voltages must satisfy to replicate experimentally-observed behaviors are listed in Table 2 in Section 2.2.

Following a similar derivation as for the simple subcircuit, we rewrite the conditions as the system of inequalities and optimization problems on coupling strength parameters given in Table 7. The inequalities in the last three rows of Table 7 involve maximizing or minimizing a quantity over the range of *f*_*Aβ*_ values. To make the APS easier to compute, we find explicit solutions to the four optimization problems in the last two rows of Table 7 using the alternative approach described above for the simple subcircuit and outlined in S1.2 Appendix and S1.3 Appendix. The solutions to the four optimization problems (Table 8, 3rd column) can be written in terms of the Lambert *W*_0_ function (4th column) with different constants *A* (5th column).

**Table 8.** Explicit solutions of *f*_*Aβ*_ for optimization problems in Table 7. The value of *f*_*Aβ*_ at extrema satisfying the four optimization problems in the last two rows of Table 7 (3rd column). All solutions (4th column) have the same form involving the Lambert *W*_0_ function with different constants *A* (5th column) (UB = upper bound, abl = ablation).

| Population | Condition | Optimization | $f_{A\beta}$ at extrema (if any) | $A$ |
| --- | --- | --- | --- | --- |
| E | UB (I1 abl) | $\max_{f_{A\beta}} \left[ \frac{g_{A\beta E}f_{A\beta}-(V_{E,max}-V_{E,rest})}{f_{I2}} \right]$ | $-\frac{\alpha_I}{2g_{A\beta I}} \left( 1 + 2A + W_0 \left( e^{2\frac{V_{I,rest}-\beta_I}{\alpha_I}} - 2A - 1 \right) \right)$ | $-\frac{V_{E,max}-V_{E,rest}}{g_{A\beta E\alpha_I}}$ |
| | Fires (I1 abl) | $\min_{f_{A\beta}} \left[ \frac{g_{A\beta E}f_{A\beta}+(V_{E,thr}-V_{E,rest})}{f_{I2}} \right]$ | | $\frac{V_{E,thr}-V_{E,rest}}{g_{A\beta E\alpha_I}}$ |
| | UB (I2 abl) | $\max_{f_{A\beta}} \left[ \frac{g_{A\beta E}f_{A\beta}-(V_{E,max}-V_{E,rest})}{f_{I1}} \right]$ | | $-\frac{V_{E,max}-V_{E,rest}}{g_{A\beta E\alpha_I}}$ |
| | Fires (I2 abl) | $\min_{f_{A\beta}} \left[ \frac{g_{A\beta E}f_{A\beta}+(V_{E,thr}-V_{E,rest})}{f_{I1}} \right]$ | | $\frac{V_{E,thr}-V_{E,rest}}{g_{A\beta E\alpha_I}}$ |

**Table 9.**
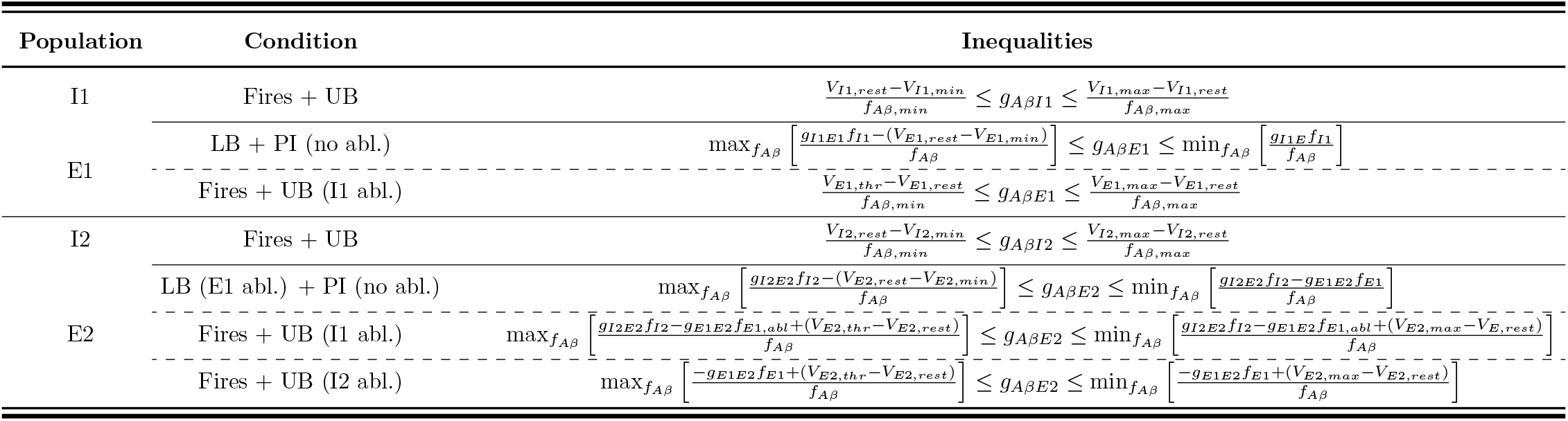
Inequalities expressed as upper and lower bounds on coupling strength values define the allowable parameter space for the dynamic subcircuit. These inequalities are obtained by algebraically manipulating the inequalities on the voltages of various populations from (Table 3) that define the APS for the dynamic subcircuit so that the inequalities are written explicitly in terms of coupling strength parameters. (UB = upper bound, LB = lower bound, abl = ablation, PI = pain inhibition)

To more easily sample from the APS, it is helpful to compute bounds on *E* population coupling strength values that are independent of other coupling strength parameters. For example, to find an upper bound on *g*_*I*1*E*_, we can use the upper bound on E during *I1* -ablation, (rewritten so the inequality is expressed as bounds on *g*_*AβE*_ rather than *g*_*I*1*E*_), along with the E lower bound given no ablations to obtain that

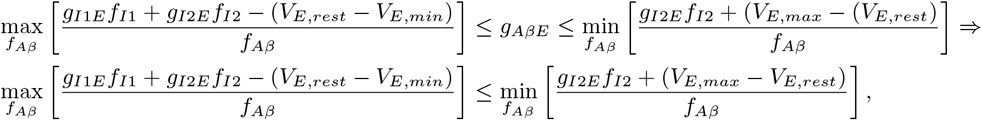

which requires in particular that for all *f*_*Aβ*_ ∈ [*f*_*Aβ*;*min*_, *f*_*Aβ*;*max*_]

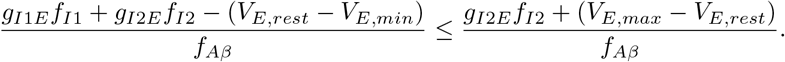

which in turn implies that

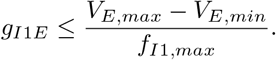

To find a lower bound on *g*_*I*1*E*_, on the other hand, we can use the E lower bound under *I1* ablation, (rewritten so the inequality is expressed as bounds on *g*_*AβE*_ rather than *g*_*I*1*E*_), along with the pain inhibition condition, i.e. that:

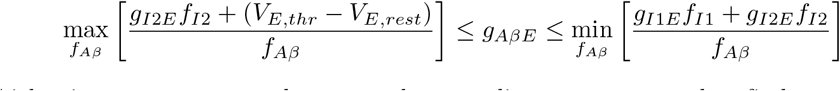

which, via an argument analogous to the preceding argument used to find an upper bound on *g*_*I*1*E*_, implies that

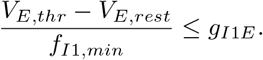

An analogous procedure produces bounds on *g*_*I*2*E*_.

#### 4.4.3 Dynamic subcircuit

We rewrite the inequalities in Table 3 in Section 2.3 as the system of inequalities and optimization problems for the coupling strength parameters given in Table 9.

We further simplify the inequalities in the second line of Table 9 by explicitly solving the optimization problems in terms of Lambert-W functions analogously to our treatment of the optimization problems in the last row of (Table 6) for the simple subcircuit. Upper and lower bounds on *g*_*I*1*E*1_ are straightforward to find, because the *I1* -*E1* portion of the dynamic subcircuit has identical constraints to those of the simple subcircuit. To find upper and lower bounds on *g*_*I*2*E*2_, we use a more computationally intensive approach. Namely, given the parameter values for the *I1* -*E1* portion of the subcircuit and given *g*_*E*1*E*2_, we find the set of *g*_*I*2*E*2_ values such that the inequalities in the bottom-most three lines of Table 9 have a solution. That is, we choose *g*_*I*2*E*2_ so that the upper bounds on *g*_*AβE*2_ are indeed larger than the lower bounds appearing in the last three lines of Table 9.

### 4.5 Normalizing the allowable parameter space (APS)

We normalize the APS so that changes in different coupling strength parameters can be compared. Since normalization requires upper and lower bounds on each coupling strength parameter, we construct a rectangular hypercube in parameter space that contains the APS. To do this, we leverage the hierarchical nature of the sets of inequalities on coupling strength parameters for each subcircuit in Tables 6, 7 and 9. The hierarchy is formed by the dependencies of inequalities on the coupling strength parameters, with inequalities higher in the hierarchy depending on parameters defined by inequalities lower in the hierarchy. Table 10 lists coupling strength parameters for each subcircuit in the order of the hierarchy formed by their inequalities (from low to high).

**Table 10.** Hierarchical order formed by the inequalities on coupling strength parameters for each subcircuit. Order of the hierarchy formed by the inequalities for coupling strength parameters shown in Tables 6, 7, and 9 listed from lowest to highest in the hierarchy.

| Subcircuit | Hierarchy of inequalities |
| --- | --- |
| Simple | $g_{A\beta I}, g_{IE}, g_{A\beta E}$ |
| Static | $g_{A\beta I1}, g_{A\beta I2}, g_{A\beta E}, g_{I1E}, g_{I2E}$ |
| Dynamic | $g_{A\beta I1}, g_{A\beta I2}, g_{I1E1}, g_{E1E2}, g_{I2E}, g_{A\beta E2}$ |

Here we describe the algorithm implemented to construct the rectangular hypercube in parameter space that contains the APS based on the hierarchy of inequalities on coupling strength parameter values. In general, consider a hierarchical set of inequalities that define bounds on elements of a parameter vector 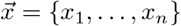 in ℝ^*n*^:

- *a*_1_ ≤ *x*_*1*_ ≤ *b*_1_
- *L*_2_(*x*_1_) ≤ *x*_2_ ≤ *U*_2_(*x*_1_) for *x*_2_ ∈ [*a*_2_; *b*_2_]
- *L*_3_(x_1_; x_2_) ≤ *x*_3_ ≤ *U*_3_(*x*_1_; *x*_2_) for *x*_3_ ∈ [*a*_3_; *b*_3_]
- *L*_*n*_(*x*_1_; *x*_2_; : : : ; *x*_*n*™1_) ≤ *x*_*n*_ ≤ *U*_*n*_(*x*_1_; *x*_2_; : : : ; *x*_*n*™1_) for *x*_*n*_ ∈ [*a*_*n*_; *b*_*n*_]

for real numbers *a*_1_ ≤ *b*_1_, … *a*_*n*_ ≤ *b*_*n*_, and real functionals *L*_1_ ≤ *U*_1_, …, *L*_*n*_ ≤ *U*_*n*_. As written, the *x*_*n*_ inequalities are the highest in the hierarchy because they depend on *x*_1_, …, *x*_*n*™1_. To identify the interval of values [*x*_*i,min*_, *x*_*i,max*_] that satisfies the inequality for each *x*_*i*_, we generate a large (at least 1000 elements), random (not necessarily uniform) sample of *_x* values satisfying the inequality system as follows:

- Uniformly at random choose a value of *x*_1_ in [*a*_1_, *b*_1_]
- Given the value of *x*_1_, uniformly at random choose a value of *x*_2_ ∈ [*L*_2_(*x*_1_), *U*_2_(*x*_1_)] if such an interval exists. If such an interval doesn’t exist, start over.
- Repeat to choose *x*_3_, …, *x*_*n*_ sequentially. If, at any step, an interval for *x*_*i*_ is not defined, start over with a new choice for *x*_1_.

We define the minimum (maximum) value of each coupling strength parameter in the APS as the minimum (maximum) value across all samples. This defines a hypercube in ℝ^*n*^ that contains the APS. To normalize, we map each element of 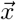 to [0, 1] as follows:

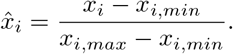

The normalized APS thus lies in the unit hypercube.

### 4.6 Sampling from the allowable parameter space (APS)

To perform analyses, we generate a uniform sampling of the APS. However, the APS is high-dimensional, can be non-convex, and is generally of an unknown shape, making it difficult to sample uniformly. In this section we briefly outline the algorithm we developed to sample the APS for each subcircuit; further details are contained in S2 Appendix.

Having found a rectangular subspace that contains the APS (Section 4.5), one approach would be to uniformly at random sample a point from the rectangular subspace, check to see if the point is in the APS, and keep it if it is. However, this naive sampling scheme is dependent on the volume of the APS in such a way that its computational complexity is exponential in *n* (see S2.2 Appendix).

#### Volume-independent sampling algorithm

Here we describe a spatially uniform sampling algorithm whose computational complexity is independent of the volume of the APS in *R*^*n*^. To do so, we sample from a cover of the normalized APS consisting of *R*^*n*^ hyperrectangles that approximates the normalized APS. Our algorithm takes the following strategy:

1. Define a set of *R*^*n*^ hyperrectangles that contains and approximates the allowable parameter space.
2. Uniformly at random select a hyperrectangle
3. Uniformly at random select a point from the hyperrectangle.
  - If the point is not in the parameter space, discard it
4. Otherwise, keep the point with probability proportional to the volume of the hyperrectangle.

Step (4) ensures that the sample is indeed uniform-in-space. Fig 8 provides an illustration of the sampling algorithm. (See S2 Appendix for details on the implementation of this algorithm, a discussion of the volume-independence of the implementation, and proof that this produces a uniform in space sampling).

**Fig 8.**
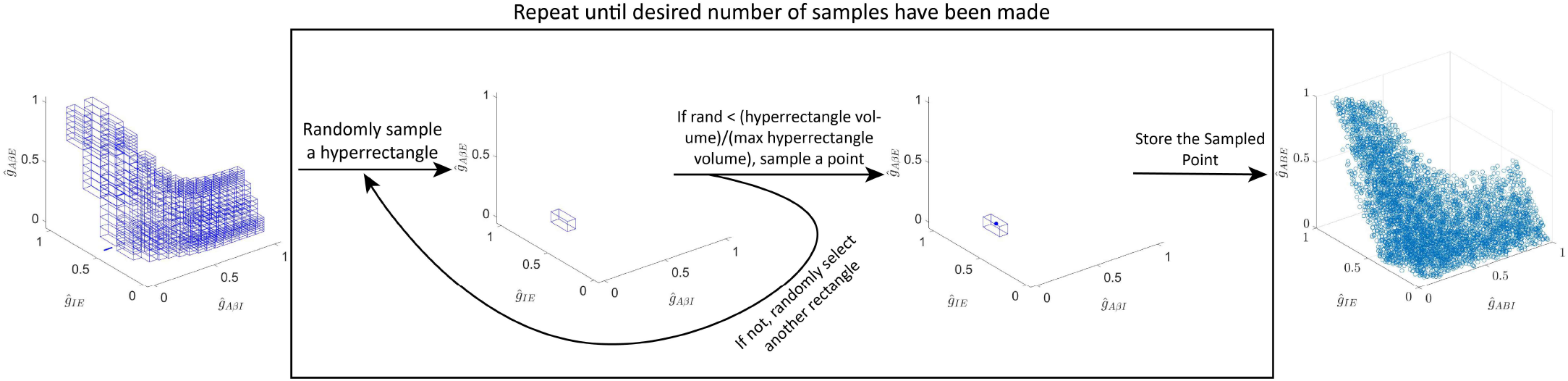
Schematic of the volume-independent sampling algorithm.

### 4.7 Defining the allodynia surface

In this section, we derive the conditions describing the allodynia surface for each subcircuit. We present the derivation in terms of a generalized circuit and conditions for a general target state to occur.

In particular, we restrict our attention to circuits where the output signal is relayed by a single neural population, which we denote by *y*. The target state is represented by the voltage of this output population *V*_*y*_ increasing above a specified threshold in response to an input signal. For our firing rate model, the steady-state average voltage *V*_*y*_ is given by

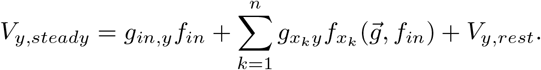

where *f*_*in*_ is the input signal and *f*_*x*_*k* are firing rates of the populations pre-synaptic to population *y*. The vector 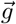 contains all the coupling strength parameters in the circuit, *g*_*ji*_ for the weight of the connection from pre-synaptic population *j* to postsynaptic population *i*. The circuit is in the target state when

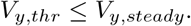

In our subcircuits, the output populations *y* are the excitatory interneuron populations that directly target the projection neurons that relay signals to the brain, namely the *E, E*, and *E*2 populations in the simple, static and dynamic subcircuits, respectively. The target state for allodynia is that the average voltage of these excitatory populations increases above the firing threshold in response to *Aβ* input. These allodynia conditions are summarized in Table 11.

**Table 11.** Allodynia conditions on each subcircuit. Allodynia conditions specifying precisely when the subcircuit is producing allodynia–when it is relaying pain-inducing stimuli in response to innocuous *f*_*Aβ*_ signals.

| Subcircuit | Conditions under which allodynia is produced |
| --- | --- |
| Simple | $V_{E,thr} \leq V_{E,steady} = g_{A\beta E}f_{A\beta} - g_{IE}f_I + V_{E,rest}$ |
| Static | $V_{E,thr} \leq V_{E,steady} = g_{A\beta E}f_{A\beta} - g_{I1E}f_{I1} - g_{I2E}f_{I2} + V_{E,rest}$ |
| Dynamic | $V_{E2,thr} \leq V_{E2,steady} = g_{A\beta E2}f_{A\beta} + g_{E1E2}f_{E1} - g_{I2E2}f_{I2} + V_{E2,rest}$ |

Using the target state condition, we can identify the coupling strength values 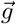 for which the target state is attainable. To do so, we rewrite the condition defining the target state as an inequality on the coupling strength *g*_*in,y*_ between the input signal and the population *y*:

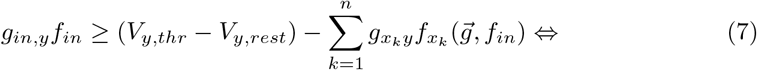

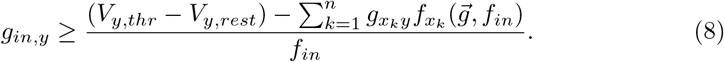

For the target state to be attainable, we don’t need to reach the target state for all values of *f*_*in*_. Instead, we can reach the target state for the value of *f*_*in*_ that minimizes the right-hand side of the preceding equation. Thus, the circuit with the set of coupling strengths 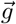 can attain the target state if and only if

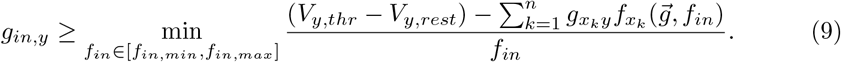

To illustrate this more concretely, the conditions on the coupling strengths for which allodynia is attainable are summarized in Table 12 for the simple, dynamic, and static subcircuits.

**Table 12.** Conditions for a subcircuit to produce allodynia. These conditions specify the sets of coupling strength values when the corresponding subcircuit will produce allodynia for some typical *f*_*Aβ*_ input.

| Subcircuit | Conditions for allodynia being possible |
| --- | --- |
| Simple | $g_{A\beta E} \geq \min_{f_{A\beta} \in [f_{A\beta,min}, f_{A\beta,max}]} \frac{g_{IE} f_I + V_{E,thr} - V_{E,rest}}{f_{A\beta}}$ |
| Static | $g_{A\beta E} \geq \min_{f_{A\beta} \in [f_{A\beta,min}, f_{A\beta,max}]} \frac{g_{I1E} f_{I1} + g_{I2E} f_{I2} + (V_{E,thr} - V_{E,rest})}{f_{A\beta}}$ |
| Dynamic | $g_{A\beta E2} \geq \min_{f_{A\beta} \in [f_{A\beta,min}, f_{A\beta,max}]} \frac{g_{I2E2} f_{I2} - g_{E1E2} f_{E1} + (V_{E,thr} - V_{E2,rest})}{f_{A\beta}}$ |

Eq (9) thus defines a target state boundary surface *S* which divides the sets of coupling strength values for which the circuit is in the target state from those for which it is not. We can express this surface as

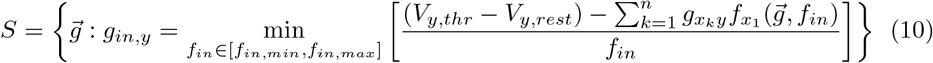

For the simple, static, and dynamic subcircuits, this boundary (Table 13) separates regions of parameter space in which the subcircuit can produce allodynia from regions where it does not.

**Table 13.** The allodynia surface for the simple, dynamic and static subcircuits, respectively. The allodynia surface separates subcircuit instantiations which can produce allodynia in response to typical *f*_*Aβ*_ signaling from those that cannot.

| Subcircuit | Allodynia surface |
| --- | --- |
| Simple | $S = \left\{ \vec{g} : g_{A\beta E} = \min_{f_{A\beta} \in [f_{A\beta,min}, f_{A\beta,max}]} \frac{g_{IE} f_I + V_{E,thr} - V_{E,rest}}{f_{A\beta}} \right\}$ |
| Static | $S_{stat} = \left\{ \vec{g} : g_{A\beta E} = \min_{f_{A\beta} \in [f_{A\beta,min}, f_{A\beta,max}]} \frac{g_{I1E} f_{I1} + g_{I2E} f_{I2} + (V_{E,thr} - V_{E,rest})}{f_{A\beta}} \right\}$ |
| Dynamic | $S_{dyn} = \left\{ \vec{g} : g_{A\beta E2} = \min_{f_{A\beta} \in [f_{A\beta,min}, f_{A\beta,max}]} \frac{g_{I2E2} f_{I2} - g_{E1E2} f_{E1} + (V_{E,thr} - V_{E2,rest})}{f_{A\beta}} \right\}$ |

### 4.8 Computing the distance between sampled points and the allodynia surface

In this section, we discuss the computational algorithm that computes the shortest path from points in the APS to the allodynia surface. Similarly as in Section 4.7, we describe the algorithm for a generalized circuit and a generalized target state boundary surface *S*.

The length of the shortest path between a point 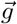 in the space of coupling strength values to the target state boundary surface *S* indicates how easy it is to move the circuit into the target state. It also identifies which coupling strengths need to change to reach the target state. We illustrate the problem of finding the shortest path to *S* by considering an arbitrary set of coupling strength values–a point in the APS which we denote 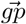. To compute the shortest path from 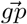 to *S*, we need to find the point 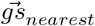 on *S* closest to 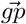. To do so, we need to solve the optimization problem:

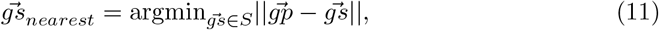

where ║·║represents the Euclidean norm.

Solving the optimization problem in Eq (11) directly would require knowing the target state surface itself or knowing important properties of it such as its gradient. However, the target state boundary surface in our work is defined via solving a minimization problem over the space of coupling strength values (excluding the coupling strength *g*_*in,y*_) and is thus difficult to include in existing optimization algorithms. Thus, we seek to solve this problem without computing an explicit representation of the target state surface *S*.

Instead, we solve the higher dimensional problem

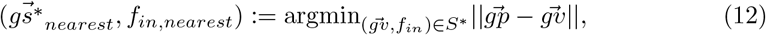

where *S*^∗^ is the following set of coupling strength-input signal pairs defined by removing the minimization from Eq (10):

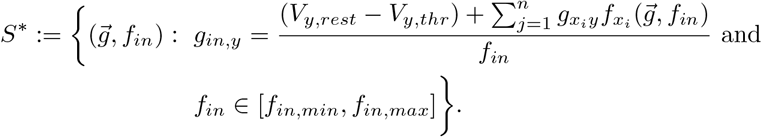

Eq (12) presents a constrained optimization problem that can be solved with high likelihood using global optimization algorithms based on stochastic gradient descent. As a result, Eq (12) is far more tractable than the original problem posed in Eq (11).

Moreover, if 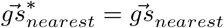, the two minimization problems are equivalent, and by solving Eq (12), we will have solved the original minimization problem Eq (11). In S3 Appendix, we describe how we solve Eq (12) using Matlab’s stochastic gradient descent-based algorithm *fmincon* in a multi-start global optimization scheme and show that if we solve Eq (12), we do indeed solve Eq (11).

### 4.9 Clustering the data based on shortest paths to the allodynia surface

We use a density-based scanning clustering algorithm coupled with data visualization to identify clusters in the APS. To do so, we work with the uniformly sampled points in the APS and cluster them according to their shortest paths to the allodynia surface.

To apply density-based clustering to the data, we use Matlab’s *dbscan* function. Briefly, *dbscan* divides the data into equivalence classes, where two datapoints are equivalent if they are sufficiently close, and identifies equivalence classes as a cluster if they contain a point which exceeds a minimum number of sufficiently close neighbors. The function takes three arguments:

1. The data to be clustered: We use the set of shortest path vectors from sampled points in the APS to the allodynia surface.
2. A sufficiently close distance *E*: we use the smallest *E* under the euclidean metric that leads to no outliers in the data.
3. The minimum number of sufficiently close neighbors to identify an equivalence class as a cluster: we take this to be 5.

Figures 3D, 5C, and 7C show parallel plots of the shortest paths to the allodynia surface for the simple, static, and dynamic subcircuits, respectively; and are colored according to the clusters assigned by density-based scanning. The shortest paths in those figures clearly divide visually into spatially-separated clusters.

## Acknowledgments

This work was funded by the National Institutes of Health through grants R01 AT010817 (AGG, SFL, BD, VB), R01 NS118769 (BD), R01 NS109170 (BD) and P20 GM103449 (JC).

## Notes

### Competing Interest Statement

The authors have declared no competing interest.

## References

1. Rikard SM, Strahan AE, Schmit KM, Guy Jr GP. Chronic Pain Among Adults—United States, 2019–2021. Morbidity and Mortality Weekly Report. 2023;72(15):379.

2. Peirs C, Williams SPG, Zhao X, Walsh CE, Gedeon JY, Cagle NE, et al. Dorsal horn circuits for persistent mechanical pain. Neuron. 2015;87(4):797–812.

3. Peirs C, Dallel R, Todd AJ. Recent advances in our understanding of the organization of dorsal horn neuron populations and their contribution to cutaneous mechanical allodynia. Journal of neural transmission. 2020;127(4):505–525.

4. Simon LS. Relieving pain in America: A blueprint for transforming prevention, care, education, and research. Journal of pain & palliative care pharmacotherapy. 2012;26(2):197–198.

5. Bear M, Connors B, Paradiso MA. Neuroscience: exploring the brain, enhanced edition: exploring the brain. Jones & Bartlett Learning; 2020.

6. Todd AJ. Neuronal circuitry for pain processing in the dorsal horn. Nature Reviews Neuroscience. 2010;11(12):823–836.

7. Lechner SG. An update on the spinal and peripheral pathways of pain signalling. e-Neuroforum. 2017;23(3):131–136.

8. Duan B, Cheng L, Ma Q. Spinal circuits transmitting mechanical pain and itch. Neuroscience bulletin. 2018;34:186–193.

9. Woolf CJ. Pain modulation in the spinal cord. Frontiers in Pain Research. 2022;3:984042.

10. Melzack R, Wall PD. Pain Mechanisms: A New Theory: A gate control system modulates sensory input from the skin before it evokes pain perception and response. Science. 1965;150(3699):971–979.

11. Braz J, Solorzano C, Wang X, Basbaum AI. Transmitting pain and itch messages: a contemporary view of the spinal cord circuits that generate gate control. Neuron. 2014;82(3):522–536.

12. Thomas Cheng H. Spinal cord mechanisms of chronic pain and clinical implications. Current pain and headache reports. 2010;14:213–220.

13. Cohen SP, Vase L, Hooten WM. Chronic pain: an update on burden, best practices, and new advances. The Lancet. 2021;397(10289):2082–2097.

14. He Y, Kim PY. Allodynia. In: StatPearls [Internet]. StatPearls Publishing; 2021.

15. Sandkuhler J. Models and mechanisms of hyperalgesia and allodynia. Physiological reviews. 2009;89(2):707–758.

16. Duan B, Cheng L, Bourane S, Britz O, Padilla C, Garcia-Campmany L, et al. Identification of spinal circuits transmitting and gating mechanical pain. Cell. 2014;159(6):1417–1432.

17. Petitjean H, Pawlowski SA, Fraine SL, Sharif B, Hamad D, Fatima T, et al. Dorsal horn parvalbumin neurons are gate-keepers of touch-evoked pain after nerve injury. Cell reports. 2015;13(6):1246–1257.

18. Cheng L, Duan B, Huang T, Zhang Y, Chen Y, Britz O, et al. Identification of spinal circuits involved in touch-evoked dynamic mechanical pain. Nature neuroscience. 2017;20(6):804–814.

19. Petitjean H, Bourojeni FB, Tsao D, Davidova A, Sotocinal SG, Mogil JS, et al. Recruitment of spinoparabrachial neurons by dorsal horn calretinin neurons. Cell reports. 2019;28(6):1429–1438.

20. Zhang Y, Liu S, Zhang YQ, Goulding M, Wang YQ, Ma Q. Timing mechanisms underlying gate control by feedforward inhibition. Neuron. 2018;99(5):941–955.

21. Hughes D, Sikander S, Kinnon C, Boyle K, Watanabe M, Callister R, et al. Morphological, neurochemical and electrophysiological features of parvalbumin-expressing cells: a likely source of axo-axonic inputs in the mouse spinal dorsal horn. The Journal of physiology. 2012;590(16):3927–3951.

22. Gobel S. Golgi studies of the neurons in layer II of the dorsal horn of the medulla (trigeminal nucleus caudalis). Journal of Comparative Neurology. 1978;180(2):395–413.

23. Freeman WJ. Mass action in the nervous system. vol. 2004. Citeseer; 1975.

24. Phillips A, Robinson PA. A quantitative model of sleep-wake dynamics based on the physiology of the brainstem ascending arousal system. Journal of Biological Rhythms. 2007;22(2):167–179.

25. Jansen BH, Rit VG. Electroencephalogram and visual evoked potential generation in a mathematical model of coupled cortical columns. Biological cybernetics. 1995;73(4):357–366.

26. Ruscheweyh R, Sandkühler J. Lamina-specific membrane and discharge properties of rat spinal dorsal horn neurones in vitro. The Journal of physiology. 2002;541(1):231–244.

27. Walcher J, Ojeda-Alonso J, Haseleu J, Oosthuizen MK, Rowe AH, Bennett NC, et al. Specialized mechanoreceptor systems in rodent glabrous skin. The Journal of Physiology. 2018;596(20):4995–5016.

28. Ester M, Kriegel HP, Sander J, Xu X. A density-based algorithm for discovering clusters in large spatial databases with noise. In: Simoudis E, Han J, Fayyad U, editors. KDD’96: Proceedings of the Second International Conference on Knowledge Discovery and Data Mining. AAAI press; 1996. p. 226–231.

29. Westlund KN. Neurophysiology of Pain: Peripheral, Spinal, Ascending, and Descending Pathways. In: Practical management of pain. Elsevier; 2022. p. 95–109.

30. Coull JA, Boudreau D, Bachand K, Prescott SA, Nault F, Sík A, et al. Trans-synaptic shift in anion gradient in spinal lamina I neurons as a mechanism of neuropathic pain. Nature. 2003;424(6951):938–942.

31. Woolf CJ, Shortland P, Coggeshall RE. Peripheral nerve injury triggers central sprouting of myelinated afferents. Nature. 1992;355(6355):75–78.

32. Moore KA, Kohno T, Karchewski LA, Scholz J, Baba H, Woolf CJ. Partial peripheral nerve injury promotes a selective loss of GABAergic inhibition in the superficial dorsal horn of the spinal cord. Journal of Neuroscience. 2002;22(15):6724–6731.

33. Medlock L, Sekiguchi K, Hong S, Dura-Bernal S, Lytton WW, Prescott SA. Multiscale computer model of the spinal dorsal horn reveals changes in network processing associated with chronic pain. Journal of Neuroscience. 2022;42(15):3133–3149.

34. Arle JE, Carlson KW, Mei L, Iftimia N, Shils JL. Mechanism of dorsal column stimulation to treat neuropathic but not nociceptive pain: analysis with a computational model. Neuromodulation: Technology at the Neural Interface. 2014;17(7):642–655.

35. Yasaka T, Tiong SY, Hughes DI, Riddell JS, Todd AJ. Populations of inhibitory and excitatory interneurons in lamina II of the adult rat spinal dorsal horn revealed by a combined electrophysiological and anatomical approach. Pain®. 2010;151(2):475–488.

36. Punnakkal P, von Schoultz C, Haenraets K, Wildner H, Zeilhofer HU. Morphological, biophysical and synaptic properties of glutamatergic neurons of the mouse spinal dorsal horn. The Journal of physiology. 2014;592(4):759–776.

37. Montbrió E, Pazó D, Roxin A. Macroscopic description for networks of spiking neurons. Phys Rev X. 2015;5(2):021028. doi:10.1103/PhysRevX.5.021028.

38. Pietras B, Pikovsky A, et al. Exact finite-dimensional description for networks of globally coupled spiking neurons. Physical Review E. 2023;107(2):024315.

39. Ferrara A, Angulo-Garcia D, Torcini A, Olmi S. Population spiking and bursting in next-generation neural masses with spike-frequency adaptation. Physical Review E. 2023;107(2):024311.

40. Chen L, Campbell SA. Exact mean-field models for spiking neural networks with adaptation. Journal of Computational Neuroscience. 2022;50(4):445–469.

41. Crodelle J, Piltz SH, Hagenauer MH, Booth V. Modeling the daily rhythm of human pain processing in the dorsal horn. PLoS Comput Biol. 2019;15(7):e1007106. doi:10.1371/journal.pcbi.1007106.

42. Peirs C, Williams SPG, Zhao X, Arokiaraj CM, Ferreira DW, Noh Mc, et al. Mechanical allodynia circuitry in the dorsal horn is defined by the nature of the injury. Neuron. 2021;109(1):73–90.

43. Inc TM. Least Squares (Model Fitting) Algorithms; 2024. Available from: https://www.mathworks.com/help/optim/ug/least-squares-model-fitting-algorithms.html.

44. Zhang TC, Janik JJ, Grill WM. Modeling effects of spinal cord stimulation on wide-dynamic range dorsal horn neurons: influence of stimulation frequency and GABAergic inhibition. Journal of neurophysiology. 2014;112(3):552–567.

45. Peyronnard J, Charron L, Lavoie J, Messier J. Motor, sympathetic and sensory innervation of rat skeletal muscles. Brain research. 1986;373(1-2):288–302.

